# StrainPanDA: linked reconstruction of strain composition and gene content profiles via pangenome-based decomposition of metagenomic data

**DOI:** 10.1101/2022.02.15.480535

**Authors:** Han Hu, Yuxiang Tan, Chenhao Li, Junyu Chen, Yan Kou, Zhenjiang Zech Xu, Yang-Yu Liu, Yan Tan, Lei Dai

## Abstract

**Background:** Microbial strains of variable functional capacities co-exist in microbiomes. Current bioinformatics methods of strain analysis cannot provide the direct linkage between strain composition and their gene contents from metagenomic data.

**Methods:** Here we present StrainPanDA (<u>Strain</u>-level <u>Pan</u>genome <u>D</u>ecomposition <u>A</u>nalysis), a novel method that uses the pangenome coverage profile of multiple metagenomic samples to simultaneously reconstruct the composition and gene content variation of co-existing strains in microbial communities.

**Results:** We systematically validate the accuracy and robustness of StrainPanDA using synthetic datasets. To demonstrate the power of gene-centric strain profiling, we then apply StrainPanDA to analyze the gut microbiome samples of infants, as well as patients treated with fecal microbiota transplantation. We show that the linked reconstruction of strain composition and gene content profiles is critical for understanding the relationship between microbial adaptation and strain-specific functions (*e.g.,* nutrient utilization, pathogenicity).

**Conclusions:** StrainPanDA can be applied to metagenomic datasets to detect association between molecular functions and microbial/host phenotypes to formulate testable hypotheses and gain novel biological insights at the strain or subspecies level.

## Introduction

There is mounting evidence that multiple within-species variants co-exist in microbiomes^1,2^. Co-existing microbial cells of the same species can have substantial variations in their gene contents (*i.e.,* accessory genome), which is largely generated by horizontal gene transfer (HGT)^3–5^. The intraspecies variation in accessory genome can lead to substantial phenotypic differences (*e.g.,* nutrient utilization, pathogenicity, antibiotic resistance) and plays an important role in microbial adaptation across environments^6–9^. Moreover, many health outcomes linked to host-associated microbiomes have been found to be consequences of the function of individual strains^8,10–14^.

Metagenomic sequencing has revolutionized microbiome studies by providing a culture-independent approach to study the composition and function of complex microbial communities. Commonly used tools for metagenomic analysis, known as metagenomics profilers, typically provide species-level taxonomic composition^15–18^. In parallel with the rapid increase of sequenced microbial isolates from culturomics studies^1,9,19,20^, high-resolution analyses of metagenomic data have revealed notable within-species variations^21,22^. Methods that enable strain-level analysis of metagenomes have been used for tracking strain transmission or dispersal^23,24^, studying population genetics of microbial strains^25^, and typing strains of specific interest^26–32^.

The gene content profile of a microbial strain determines its biological function. To date, the majority of strain-level analysis methods use single nucleotide variants (SNVs) to identify strain composition^25,28,29,33,34^. By assuming an association between SNV haplotypes and gene content profiles^4^, SNV-based methods can indirectly profile the within-species gene content variation. However, for many species, it has been shown that SNV haplotypes cannot capture microbial genetic diversification resulting from HGT^4^. Alternatively, current pangenome-based method can infer the gene content of the dominant strain in a metagenomic sample^35^, but fails to provide the abundance and gene contents of co-existing strains within the sample. Establishing the linkage of composition and gene contents of co-existing within-species variants can provide crucial insights into microbial adaptation and microbiome-host interactions, but is inaccessible from currently available reference-based bioinformatics tools.

To meet the increasing needs of strain-level functional inference from metagenomics data, we developed a novel method StrainPanDA (<u>Strain</u>-level <u>Pang</u>enome <u>D</u>ecomposition <u>A</u>nalysis) to simultaneously reconstruct the composition and gene contents of co-existing strains using the pangenome coverage profile from metagenomic data (**Fig. 1**). We validated the performance of StrainPanDA with a comprehensive collection of synthetic datasets and showed that StrainPanDA was able to accurately infer strain composition and gene content profiles from metagenomic data. To demonstrate the practical use of StrainPanDA in human microbiome studies, we analyzed longitudinal gut microbiome samples of mother-infant pairs^36^ and patients treated with fecal microbiota transplantation (FMT)^29,37^. We found that StrainPanDA was able to identify association between strain-specific functions and microbial adaptation (or host phenotypes), leading to novel biological insights at the infraspecific level and testable hypotheses of molecular mechanisms.

**Figure 1.**
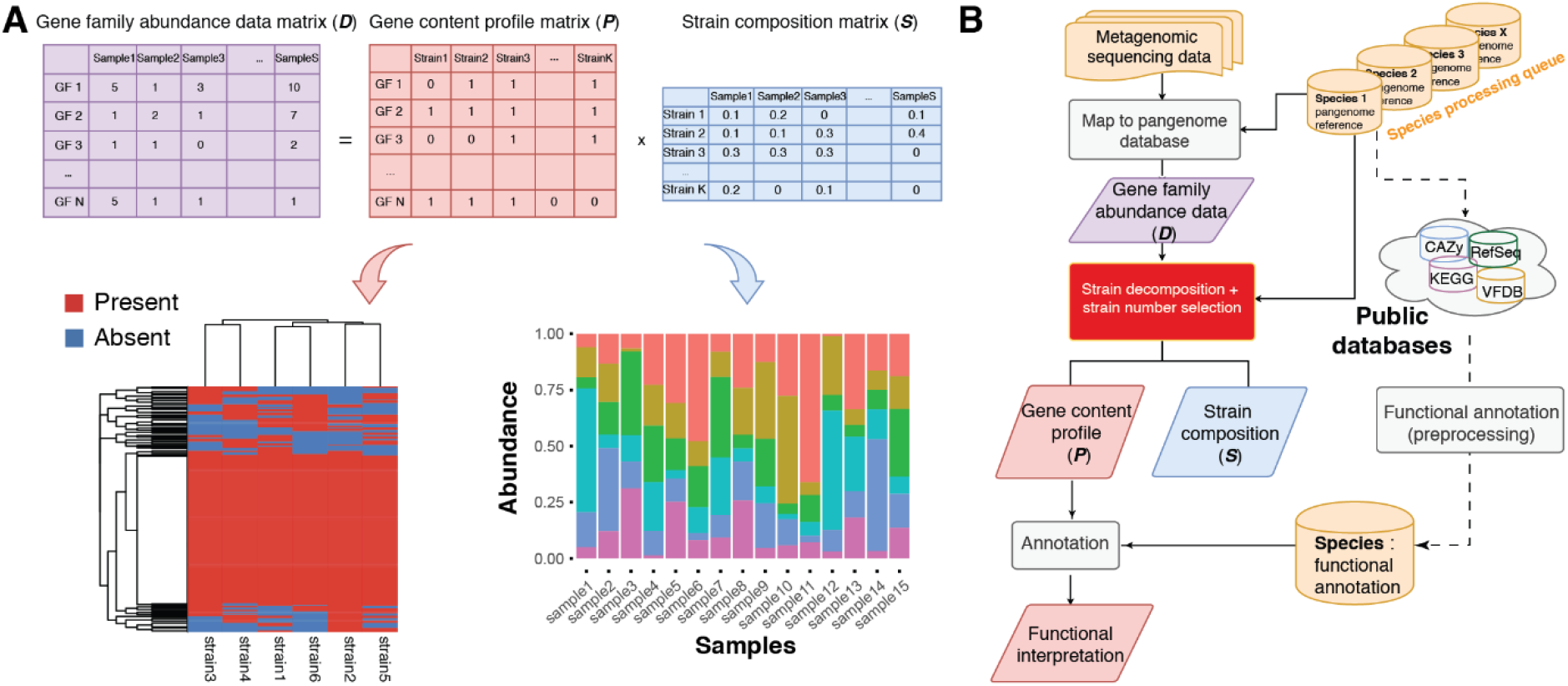
Illustration of the StrainPanDA workflow. (**A**) The gene family abundance data matrix ***D*** is decomposed into the product of two matrices ***P*** and ***S*** via non-negative matrix factorization (**Methods**). The gene content profile matrix ***P*** is a binary matrix that denotes the presence/absence of gene families in each strain. The strain composition matrix ***S*** represents the relative abundance of co-existing strains in each sample. In the illustrated example, the size of metagenomic samples S = 15, the number of strains K = 6. GF: Gene Family. (**B**) The workflow of StrainPanDA analysis is performed in a species-by-species manner, including mapping metagenomic reads to the pangenome database, strain decomposition and functional annotation of gene family profiles.

## Results

### Decomposition of the pangenome coverage profile to infer strain composition and gene content

The pangenome coverage profile of a microbial species from metagenomic data is composed of the gene contents of all co-existing strains. If there are multiple metagenomic samples with varying strain compositions, in principle it is possible to infer the composition of strains within the sample as well as the gene contents of each strain from the pangenome coverage profile^38^. Building on this intuition, the main algorithm of StrainPanDA aims to decompose the gene family abundance data matrix ***D*** into the product of two matrices, the gene content profile matrix ***P*** and the strain composition matrix ***S*** (**Fig. 1A,** see **Methods** for details). Here the pangenome coverage profile from metagenomic data is represented by matrix ***D***, where *D_ij_* is the normalized count of gene family *i* in metagenomic sample *j.* The gene contents of co-existing strains are represented by binary matrix *P,* where the element *P_ij_* indicates the presence/absence of gene family *i* in strain *j.* The composition of co-existing strains across samples is represented by *S,* where *S_ij_* is the relative abundance of strain *i* in sample *j* (*S_ij_* ≥ 0 and ∑_*i*_ *S_ij_* = 1). In the implementation of StrainPanDA, the gene family abundance matrix ***D*** is decomposed by non-negative matrix factorization (NMF) ^39–41^ to solve for matrices ***P*** and ***S***. This processing allows StrainPanDA to simultaneously delineate the composition and gene contents variation of co-existing strains.

The StrainPanDA software provides a fully automated workflow of strain analysis (**Fig. 1B**), which imports raw sequencing data from multiple metagenomic samples, performs reads mapping, strain decomposition, and downstream annotations (see **Methods** for details). To assist the interpretation of gene content variation in strains, StrainPanDA incorporates functional annotation from several databases, including but not limited to KEGG^42^, CAZy^43^, and VFDB^44^.

### StrainPanDA provides accurate predictions of strain composition and gene family profiles in synthetic data

We validated the performance of StrainPanDA using synthetic metagenomic data (**Methods**). For synthetic mixtures of *E. coli* strains (ranging from 2 to 8 strains, see **Methods**), the strain composition predicted by StrainPanDA was close to the actual composition (Ground Truth) (**Fig. 2A**). For quantitative comparison, we calculated the Jensen-Shannon divergence (JSD) and Matthews Correlation Coefficient (MCC) between the predicted and actual strain composition of simulated samples. JSD and MCC have been widely used in the evaluation of strain analysis tools ^28,30,33^. The predicted strain composition by StrainPanDA was better than or similar to the state-of-the-art SNV-based methods, including StrainEst^30^ and PStrain^33^ (the latter was modified based on ConStrains^28^) (**Fig. 2B, Supplementary Fig. 4**). Furthermore, we generated synthetic mixtures of *E. coli* strains with varying levels of sequencing errors (**Supplementary Fig. 2**), sequencing depths (**Supplementary Fig. 3**) and different background noise (mixed with different metagenomic datasets, **Supplementary Table 1, Supplementary Fig. 4**). In comparison to SNV-based methods, the performance of StrainPanDA in predicting strain composition was robust.

**Figure 2.**
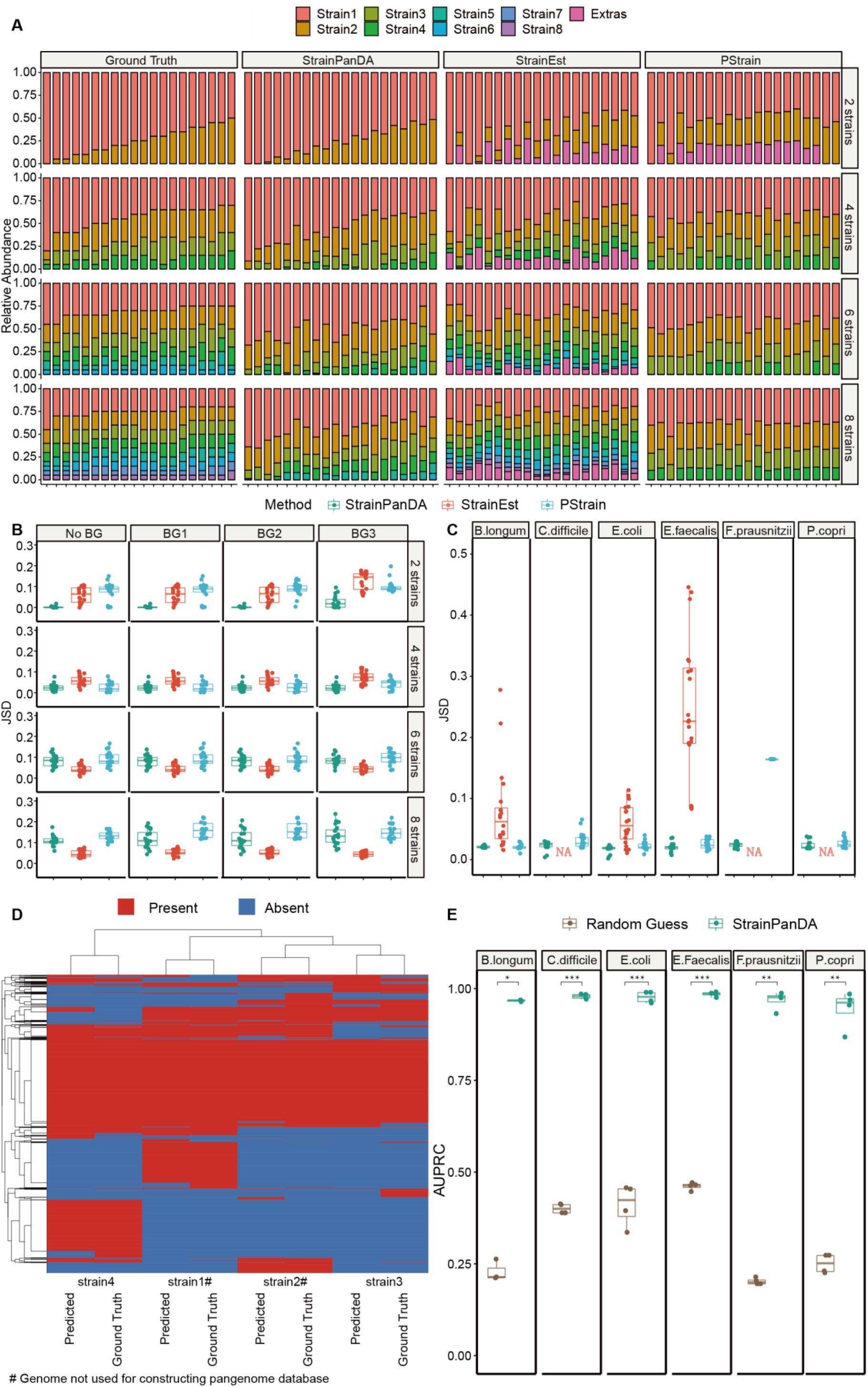
Validation of StrainPanDA using synthetic metagenomic data. (**A**) Comparison between the actual strain composition (Ground Truth) and the strain composition predicted by StrainPanDA and existing tools (StrainEst and PStrain) in synthetic mixtures of *E. coli* strains (pWGS dataset, 1 × sequencing depth, see **Methods**). The number of actual *E. coli* strains in the mixture (n = 2, 4, 6 and 8) is shown in rows. Each stacked bar is one simulated sample. Strains are displayed by the order of sorted relative abundance. If the number of predicted strains exceeds the number of actual strains, the extra strains are grouped into “Extras”. (**B**) Jensen-Shannon Divergence (JSD) between the actual and predicted strain composition. No BG: No Background; BG1/BG2/BG3: synthetic data of *E. coli* strains mixed with three different metagenomic datasets as background (WGSBG dataset, 100-fold background, see **Methods**). Each dot represents one simulated sample (n = 20). (**C**) JSD between the actual and predicted strain composition is evaluated for different microbial species. Each dot represents one simulated sample (n = 24). Outputs not available are marked as “NA”. (**D**) The actual and predicted gene family profiles of *E. coli* strains (the synthetic data used is the same as panel **A**). Each row is one gene family, and each column is one strain. Hierarchical clustering is based on Euclidean distance. (**E**) The area under the Precision-Recall Curve (AUPRC) for the gene family profiles of co-existing strains is evaluated for different microbial species. Each dot represents the AUPRC of one strain (n = 4 strains). Brown dots represent random guesses of gene family profiles (see **Methods**). P values from t-test: *P < 0.05, ***P < 0.001, ****P < 0.0001.

To evaluate the performance of StrainPanDA in different bacterial species, we generated synthetic data for common human gut bacterial species (*Bifidobacterium longum, Clostridium difficile, Enterococcus faecalis, Faecalibacterium prausnitzii* and *Prevotella copri*) (**Supplementary Table 2**, see **Methods**). In comparison to other methods, StrainPanDA made the most accurate prediction of strain composition across all species (with JSD as 0.021 ± 0.006) (**Fig. 2C**).

While current SNV-based methods can reconstruct the composition of co-existing strains from metagenomic samples, they could not directly provide the gene contents of the predicted strains. Here we show that StrainPanDA allows simultaneous reconstruction of strain composition in each metagenomic sample and the gene content variations among strains. In synthetic mixtures of *E. coli* strains, the predicted gene family profiles by StrainPanDA were close to the actual profiles (**Fig. 2D, Supplementary Fig. 5;** precision=0.91-0.96, recall=0.87-0.96, for the 4 strains in synthetic mixture). In particular, we note that StrainPanDA is able to infer the gene family profile of strains not included in the pre-built reference genome database. The area under the Precision-Recall Curve (AUPRC) was over 0.95 for all *E. coli* strains, indicating that StrainPanDA was able to reconstruct the gene contents of microbial strains with high sensitivity and precision (**Supplementary Fig. 6**).

We further evaluated the predicted gene family profiles of the human gut bacterial species included in the synthetic data. The AUPRC was on average above 0.9 and significantly better than random guesses (**Fig. 2E**). The predicted gene family profiles were robust to sequencing errors, sequencing depths and the background of real metagenomic data (**Supplementary Table 4**). Moreover, to demonstrate the ability of StrainPanDA to identify strain-specific genes, the pathogenic *E. coli* outbreak strain O104^45^ was introduced in a synthetic mixture with other *E. coli* strains (**Supplementary Table 5**). All outbreak-related gene families were successfully recovered by StrainPanDA (**Supplementary Fig. 7**). Finally, the predicted gene family profiles for the dominant strain from StrainPanDA were overall more similar to the ground truth compared to the predictions from PanPhlAn^35^, which is a pangenome based method that can only profile the gene content of the dominant strain in each sample (**Supplementary Fig. 8**).

Taken together, our benchmarking results demonstrate that StrainPanDA provides accurate predictions of compositional profiles and gene contents of co-existing strains from metagenomic samples. In the following sections, we will demonstrate the application of StrainPanDA in two longitudinal metagenomic datasets to elucidate the diversity of gut microbiome at the subspecies level.

### Succession of *B. longum* subspecies in infant gut microbiome is associated with breastfeeding patterns and the selection on nutrient utilization

The direct inference of both the population structure and gene content variation at the strain-level is crucial to understanding the ecology and evolution of microbial communities. Here we apply StrainPanDA to study the adaptation of co-existing bacterial subspecies in the infant gut microbiomes. We analyzed a previously published dataset that includes gut metagenomic samples from ~100 mother-infant pairs (infants were sampled at three time points: newborn, 4-month and 12-month)^36^ (**Supplementary Table 6**). At the species level, the authors found that the composition of infant gut microbiome had distinctive features at each sampled time point, and the cessation of breastfeeding was clearly associated with the maturation of an infant gut microbiome into an adult-like microbiome^36^.

We focused on the infraspecific analysis of *Bifidobacterium longum*, which is known to play an important role in the development of the infant gut microbiome^36,46–50^ and was found to be enriched in 4-month infant samples in this study (**Supplementary Fig. 9**). Interestingly, we discovered a clear pattern of succession in the subspecies composition of *B. longum* over time, *i.e.,* a shift in the dominant subspecies (**Fig. 3A**). Among the three *B. longum* subspecies predicted by StrainPanDA, *B. longum* subspecies 2 was dominant in the gut microbiomes of mothers. In the gut microbiomes of infants, *B. longum* subspecies 3 was most prevalent for newborns, while subspecies 1 transiently increased at the intermediate time point (at 4-months) and then was outcompeted by subspecies 2 (at 12-months). Based on the diet history provided in the original study, we further grouped infant samples into two different categories, “discontinued breastfeeding” and “continued breastfeeding” between successive timepoints (**Methods**). We found clear evidence that the relative abundance of *B. longum* subspecies 1 was enriched in infants that continued breastfeeding (**Fig. 3B**). By contrast, once breastfeeding was discontinued, *B. longum* subspecies 1 was taken over by subspecies 2.

**Figure 3.**
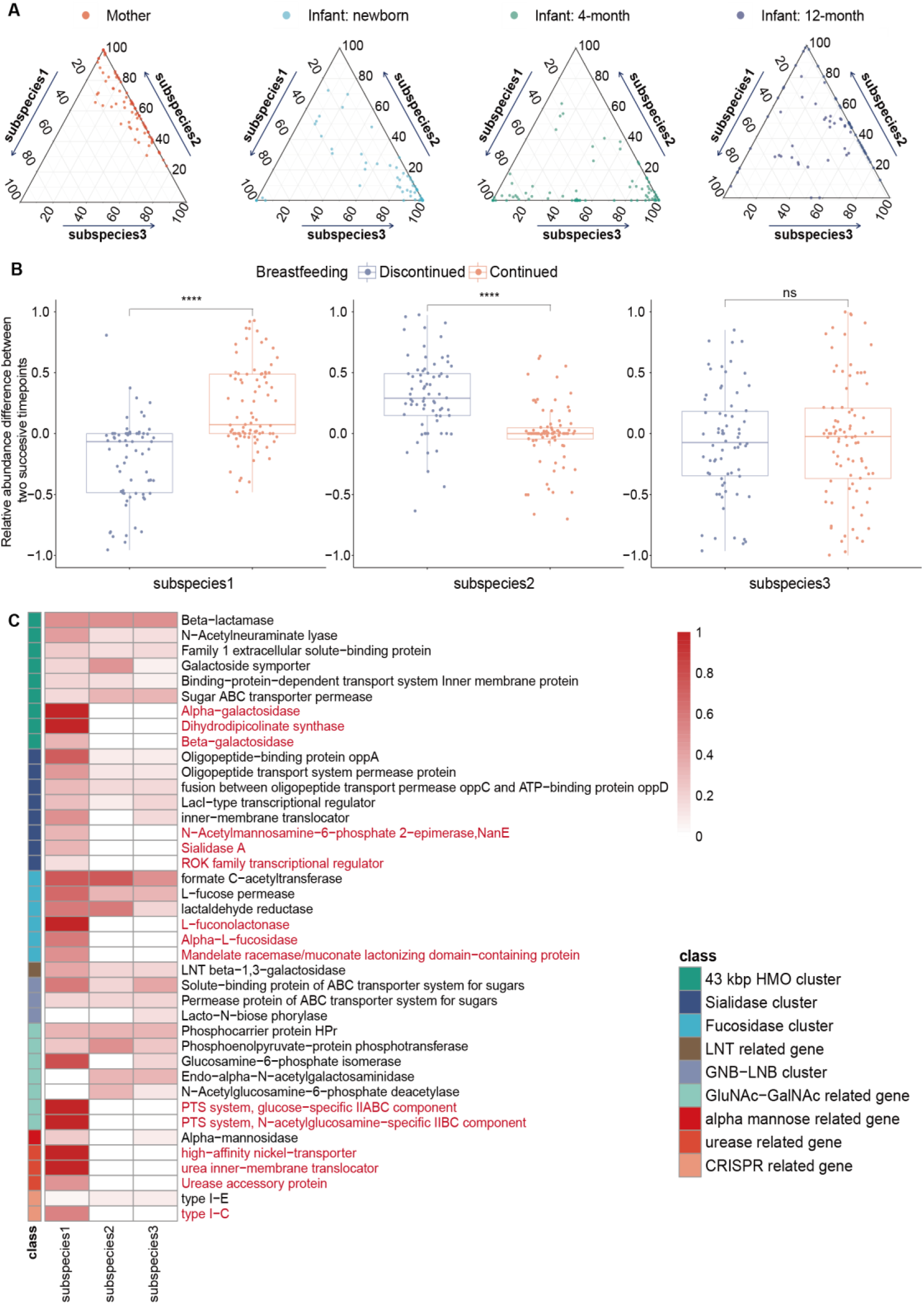
Succession of *B. longum* subspecies in infant gut microbiome can be attributed to the selection on nutrient utilization. (**A**) Ternary plots of the predicted composition of three subspecies of B. *longum* from mothers and infants of multiple time points (newborns, 4-month, 12-month). Each dot represents one sample. **(B)** The shift in the relative abundance of B. *longum* subspecies between successive time points. According to the breastfeeding status at the subsequent time point, infants are divided into two groups (purple: discontinued breastfeeding, N=86; red: continued breastfeeding, N=71). ****: p-value <0.00005, ns: not significant, Student’s t-test. (**C**) Gene family profiles of predicted *B. longum* subspecies. Gene families related to the metabolism of host glycans, urease and CRISPR (KEGG annotations) are selected for display. The color bar on the left indicates the class of gene clusters. Each row is a subclass of gene families and unique gene families of subspecies 1 are marked in red. The color scale in heatmap indicates the normalized gene family coverage in the specific subclass (*i.e.*, the fraction of detected gene families belonging to the subclass).

The association between *B. longum* subspecies composition and breastfeeding patterns suggests natural selection on nutrient utilization functions (**Fig. 3C**). Based on functional annotations of KEGG^42^ and CAZy^43^, we found clear variations in nutrient utilization genes among the predicted *B. longum* subspecies. *B. longum* subspecies 1 had unique gene families (marked in red, **Fig. 3C**) that are key enzymes related to human milk oligosaccharide (HMO), including galactosidase, alpha-L-fucosidase, sialidase and their corresponding CAZy groups (GH33, GH29 and GH95) (**Supplementary Fig. 12-13** and **Supplementary Table 7**). In addition, the urease-related gene families were only found in the gene family profile of subspecies 1. Therefore, the unique functional potential of *B. longum* subspecies 1 in utilizing HMO and urea^51^ from breast milk could confer a fitness advantage under breastfeeding, consistent with our observations of its transient dominance at 4 months in infant gut metagenomes.

Our finding is consistent with previous reports^52^ on HMO utilization genes in some *B. longum* strains as well as observed changes in the frequency of *B. longum* subspecies *infantis* after weaning^4,46,47,53,54^. The functional profile of subspecies 1, in comparison to *B. longum* reference genomes, suggests that it may correspond to the *B. longum* subsp*. infantis* (**Supplementary Fig. 14**), which has been isolated from infants and known to be associated with breastfeeding^46,55,56^.

Overall, we show that StrainPanDA is able to identify associations between strain-specific functions (via reconstruction of gene contents) and adaptation (via reconstruction of strain composition), leading to novel biological insights and testable hypotheses about microbial evolution at the subspecies level.

### Analysis of post-FMT gut metagenomes reveals individualized subspecies profiles and subspecies-specific functions

Fecal microbiota transplantation (FMT) introduces bacterial strains from healthy donors into recipients and has profound impacts on the structure and function of the recipient’s gut microbiota^57,58^. Here we apply StrainPanDA to analyze the metagenomic samples from FMT recipients in a clinical trial to treat Crohn’s disease^37^, including 17 patients (8 in the FMT group and 9 in the sham group) and multiple samples (4-8 time points) for each patient. We analyzed the subspecies composition of commonly observed bacterial species in the human gut metagenome (**Supplementary Table 8)**. Hierarchical clustering of the predicted subspecies compositional profiles revealed strong individual signatures, which remained stable throughout 24 weeks (**Fig. 4A**). The pairwise distance in subspecies composition profiles among samples of the same individual (sampled at multiple time points, *i.e.*, the “intrasubject” group) was significantly lower than the pairwise distance among samples of different individuals (*i.e.*, the “intersubject” group) (**Fig. 4B**, p-value < 10^−15^), similar to the pattern at the species level. Furthermore, we found that the dissimilarity in subspecies composition between paired FMT donors and recipients (*i.e.,* the “donor-recipient” group) was significantly lower than the “intersubject” group (p-value < 0.001), indicating the engraftment of donor strains and co-existence of donor and recipient strains^37^. We noted that the engraftment of donor gut bacteria was more obvious at the subspecies level (effect size=1.4) than at the species level (effect size=0.78) (**Fig. 4B**). Similarly, we applied StrainPanDA to analyze an independent FMT dataset of metagenomic samples from patients with *Clostridium difficile* infection^29^. We observed a clear pattern of subspecies engraftment in post-FMT gut metagenomes, consistent with SNV-based strain analysis in the original study^29^ (**Supplementary Fig. 15**). Overall, we show that StrainPanDA is able to delineate the difference in subspecies composition among individuals and track the transmission of strains.

**Figure 4.**
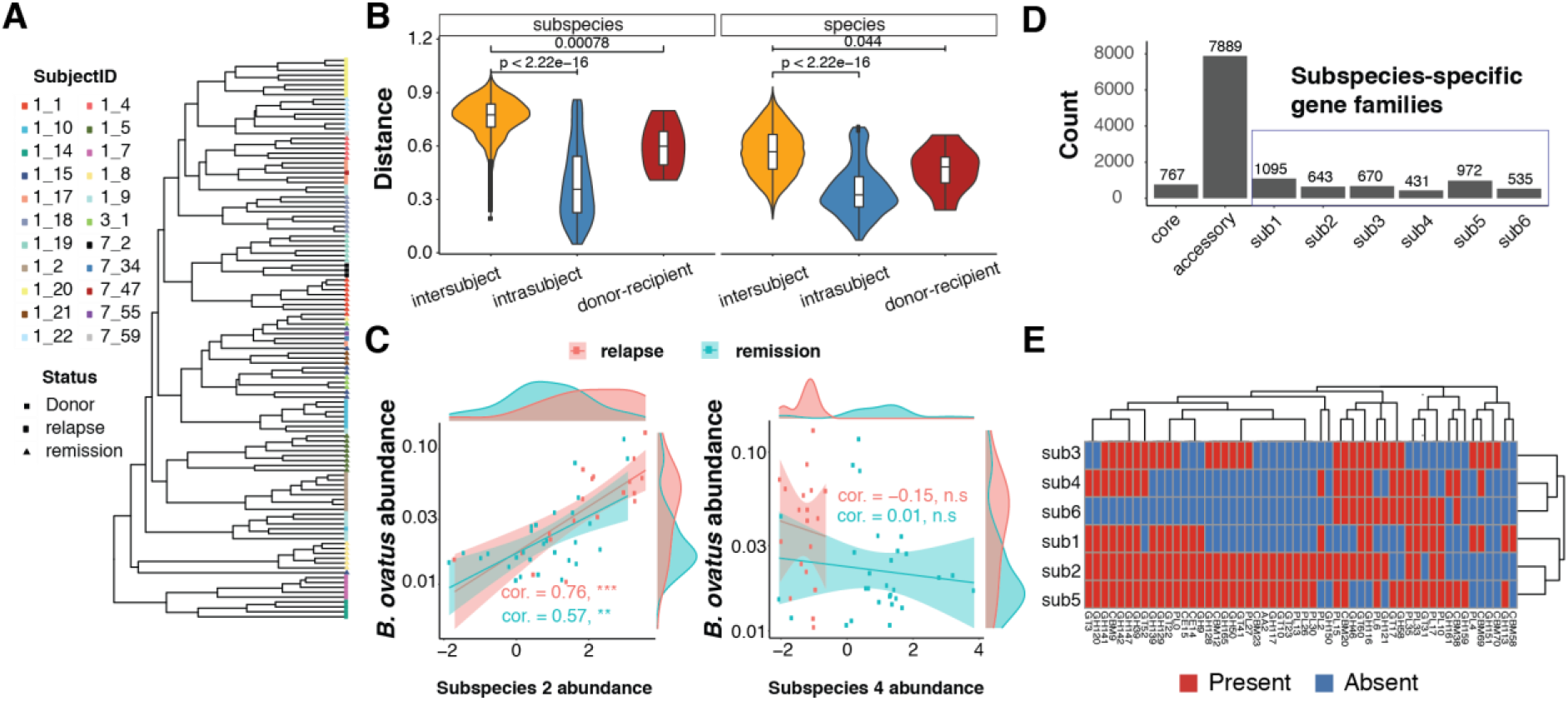
Analysis of post-FMT gut metagenomes reveals individualized subspecies profiles and the association between subspecies-specific functions and phenotypes. (**A**) Hierarchical clustering of predicted subspecies compositional profiles of common gut species (**Supplementary Table 8**) reveals strong individual signatures. The subject IDs were collected from the original paper (Kong *et al.* 2020) and marked by different colors. (**B**) The dissimilarity (Bray-Curtis dissimilarity) in subspecies composition and species composition between samples. The pairs are classified into three groups for comparison: intersubject (samples from different individuals), intrasubject (samples from the same individuals) and donor-recipient (FMT donor vs. the post-FMT sample of the recipient). (**C**) The relationship between the relative abundance of *Bacteroides ovatus* and its subspecies (normalized by centered log-ratio transformation). Lines represent fitted linear regression (shaded areas: 95% confidence interval). The density plots on the side shows the distribution of the corresponding variables. At the species level, *Bacteroides ovatus* is enriched in the relapse group. (**D**) The summary of pangenome information of predicted *B. ovatus* subspecies. (**E**) Gene family profiles of Carbohydrate-Active enZYmes (CAZy) in predicted *B. ovatus* subspecies. CAZy genes shared by all the subspecies are not shown.

To elucidate the potential role of gut microbiome in the maintenance of remission in Crohn’s disease patients, we further investigated the strain-level genetic signatures associated with post-FMT clinical outcomes. The original study has noted the enrichment of Bacteroidetes species in patients relapsed after FMT^37^. We focused our analysis on *Bacteroides ovatus,* which was found to be enriched in relapsed individuals (FDR-adjusted p-value = 0.2) (**Fig. 4C-E**). Among the predicted subspecies of *B. ovatus*, the relative abundance of subspecies 2 was positively correlated with the abundance of species-level *B. ovatus* in gut metagenomes (Spearman correlation = 0.64, and FDR-adjusted p-value < 10^−6^, **Supplementary Fig. 16**). We found substantial gene content variation among different *B. ovatus* subspecies (**Fig. 4D**). Interestingly, we found that *B. ovatus* subspecies 2 had more CAZy family genes than others, indicating its functional potential to utilize diverse carbon sources and potential fitness advantages (**Fig. 4E**). Thus, the strain-specific metabolic functions of *B. ovatus* subspecies 2 may explain its dominance within the species (~20%) as well as its positive correlation with species abundance (**Fig. 4C** and **Supplementary Fig. 16**). In addition, *B. ovatus* subspecies 2 carried several strain-specific virulence factor genes (*e.g.,* type IV secretion system, cholesterol-dependent cytolysin), which may contribute to the positive association between *B. ovatus* and post-FMT relapse (**Supplementary Fig. 16**). For example, cholesterol-dependent cytolysin is a pore forming toxin that can disrupt the host plasma membrane^59^, whose integrity has been linked to inflammatory bowel disease^60^. In addition, we noted that *B. ovatus* subspecies 4 was more abundant in the remission group (**Fig. 4C**, FDR-adjusted p-value < 10^−5^); whether this *B. ovatus* subspecies contributes to post-FMT remission remains to be validated in future studies. Similarly, we performed StrainPanDA analysis for *Bacteroides vulgatus*, which was also enriched in relapsed individuals, and found clear functional variation among its subspecies (**Supplementary Fig. 17**).

In summary, we show that the linkage of strain composition and gene contents provided by StrainPanDA can greatly facilitate our understanding of microbial ecology and evolution beyond the species level. For microbes closely related to host health, this linkage helps formulate testable hypotheses on the association between molecular functions (*e.g.*, pathogenicity) and clinical outcomes, which can be directly tested in experiments of isolated microbial strains.

## Discussion

Here we report a novel method, StrainPanDA (<u>Strain</u>-level <u>Pan</u>genome <u>D</u>ecomposition <u>A</u>nalysis), to simultaneously profile the composition of co-existing strains and their corresponding gene content from metagenomics data. Our benchmarking results showed that StrainPanDA provided accurate and robust predictions from synthetic data. The predicted strain composition was better than or comparable to state-of-the-art methods; meanwhile, the predicted gene content profile was close to the actual profile, even for strains not included in the pre-built reference genome database. Furthermore, we applied StrainPanDA to metagenomic datasets to resolve within-species variation of bacterial taxa of interest. For example, we found that the composition of *B. longum* subspecies in infant gut microbiomes was associated with dietary shifts and the unique functional potential of certain *B. longum* subspecies in utilizing nutrients from breast milk might confer a fitness advantage. We demonstrated that the linkage of strain abundance and gene contents could lead to direct functional interpretations and testable hypothesis.

To study within-species gene content variation, current SNV-based methods implicitly assume the association between SNV haplotypes and gene content. However, many microbial genomes with high similarity in core genome have less than 70% of genes in common^3^, indicating that the indirect inference of gene content by SNV-based methods may be insufficient. In contrast, StrainPanDA adopts the pangenome-based approach to directly infer the gene content of multiple co-existing within-species variants. With increasing availability of sequenced microbial isolates from environmental and host-associated microbiomes, we expect the pangenome database to expand accordingly. The prediction of StrainPanDA relies on the pangenome database, but it is not limited to the profiles of available reference genomes; thus, StrainPanDA can also be used to identify novel strains, as long as the relevant gene families are included in the pangenome. Although we focused the comparison to reference-based methods, it is worth noting that complementary approaches based on metagenome-assembled genomes (MAGs) such as DESMAN^61,62,63^ can also identify novel strains from metagenomic data. However, the MAG-based methods require much higher sequencing depth than reference-based methods, which prohibits their application to species with low abundance. In comparison, sequencing depth is not a limiting factor for StrainPanDA (**Supplementary Fig. 3-4**).

StrainPanDA is designed to fit the analysis of multiple metagenomic samples with shared within-species variants, such as longitudinal studies. The performance of StrainPanDA, including the accuracy of predicted strain composition and gene content profiles, improves with sample size (**Supplementary Fig. 18**) and sequencing depth (**Supplementary Fig. 3**). To apply StrainPanDA on a typical metagenomic dataset, it would be desirable to have at least 10 samples and the relative abundance of the species of interest to be above 1%. Due to the nature of StrainPanDA’s algorithm, it may be difficult to disentangle within-species variants with highly similar gene content profiles, thus StrainPanDA is most suitable for analysis at the level of subspecies^3^. Finally, in comparison to MAG-based methods, StrainPanDA has minimal requirements for computing resources (about 400 minutes on a 20-sample dataset for one species) and can be scaled to process multiple species in parallel.

## Conclusions

In summary, we show that StrainPanDA is able to provide accurate profiling of strain composition and gene content from metagenomic data. We envision that the application of StrainPanDA to the rapidly increasing metagenomic datasets, especially in the context of spatio-temporal characterization of microbiomes^64–67^, will help elucidate novel associations between molecular functions and microbial/host phenotypes as well as microbial ecology and evolution at the infraspecies level.

## Materials and methods

### Generation of pangenome database and mapping of metagenomic data

Pangenome database of bacterial species analyzed in this study was created following the steps recommended by PanPhlAn (version 1.2.8)^35^. For each bacterial species, genomes were downloaded from NCBI. Average Nucleotide Identity (ANI) between genomes was calculated by mash (version 1.1)^68^. Representative strains (pairwise ANI ≤ 99%) were selected and used as reference genomes for pangenome construction. The annotated genes were extracted from the reference genomes and clustered into gene families at 95% identity by usearch (v7) ^69^ to create the pangenome database. Shotgun metagenomic data was mapped to the pangenome database by PanPhlAn^35^ (version 1.2.8), which used Bowtie2 (version 2.4.1)^70^ and SAMtools (version 0.1.19)^71^ to map and count the reads, respectively. A gene family profile was generated by summing up the read counts of genes (normalized by RPKM) belonging to the same gene family. The gene family profiles of all metagenomic samples were grouped into a single gene family profile matrix. To account for potential noise in reads mapping, the gene family abundance was trimmed to 0 if RPKM value was below the cutoff (10, by default). After trimming, gene families absent in all samples were removed from further analysis. In addition, samples were filtered out if the number of gene families detected was below 0.9 ×g_*min*_ (g_*min*_ is the minimum number of gene families found in all reference genomes).

### StrainPanDA algorithm

The core algorithm of StrainPanDA decomposes the gene family abundance data matrix (*D*) of the microbial species of interest into the product of two matrices (**Fig. 1**):

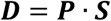

The gene family abundance data matrix *D* is a N×S non-negative matrix, where *D_ij_* is the normalized count of gene family *i* in metagenomic sample *j.* The gene content profile matrix *P* is a N×K binary matrix, where *P_ij_* is 1 if the gene family *i* is present in strain *j* and 0 otherwise. The strain composition matrix *S* is a K×S matrix, where *S_ij_* is the relative abundance of strain *i* in sample *j* (*S_ij_* ≥ 0 and ∑_*i*_ *S_ij_* = 1). N is the number of gene families in the pangenome of the microbial species of interest, S is the number of metagenomic samples, K is the number of strains (*i.e.,* factorization rank).

To estimate ***P*** and ***S***, we approximate the solution ***P’*** and ***S’*** using non-negative matrix factorization (NMF), considering the non-negative constraints on both matrices (optimized using the “snmf/r” algorithm implemented in the R package ‘NMF’^40,41^, version 0.21.0). The addition of sparsity constraints (*i.e.*, regularization terms in the objective function) ensures uniqueness of factorization^41,72^. The ***S’*** matrix is then scaled into relative abundances. We binarize the approximated ***P’*** matrix, following the assumption that the matrix elements corresponding to “present” gene families should have higher values than “absent” gene families, and the matrix elements should have a tight distribution due to the expectation that ***P*** is a binary matrix (see **Supplementary Fig. 1B**). Briefly, we find the peak of the probabilistic density curve (*p_max_*) for each strain *j*, where the number of matrix elements on the right of the peak (*P_ij_* > *p_max_*) is equal to the expected number of gene families of the species of interest (*i.e.*, averaged over all reference genomes in the pangenome database). We then cut the density curve at *θ* between the selected peak and 0 (*θ* = 0.5 × *p_max_*, by default), where the gene families with a weight greater than *θ* are considered as present. The confidence score *C_ij_* for gene family *i* in sample *j* was assigned to every gene family:

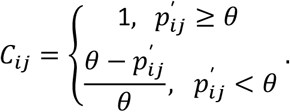

The confidence scores were used to rank gene presence predictions for generating the Precision-Recall curves in the benchmarking experiments.

To select the proper number of strains (*i.e.*, the rank of NMF), we parsimoniously select the least number of strains from a range of 1 to 12 (by default) satisfying the following criteria: 1) The mean relative abundance across all the samples of any strain should be greater than τ_2_ (τ_2_ = 0.1 by default), 2) the number of gene families of all strains should be greater than τ_3_× *g*_min_ (τ_3_ = 0.5 by default), *g*_min_ is the minimum number of gene families found in all reference genomes, and 3) the gene family profiles between a pair of strains should have Jaccard distance larger than τ_1_ (τ_1_ = 0.1 by default). The program also provides an option to accept user-specified number of strains set. In this study, we did not set the number of strains a priori in benchmarking and applications of StrainPanDA.

### Benchmarking StrainPanDA with synthetic data

#### Synthetic data of E. coli strains

We generated four types of simulated sequencing reads: 1) Error-Free (ErrFree): pick random fragments from the reference genome of *E. coli* by ART simulator^73^ (version 2016.06.05; parameter: -ef -ss HS25 -l 150 -m 270 -s 27). 2) ART with sequencing errors (ARTErr): use ART to add sequencing errors on top of ErrFree reads (parameter: -ss HS25 -l 150 -m 270 -s 27). 3) pure WGS (pWGS): randomly draw reads from the whole genome sequencing data of selected strains by seq-tk (https://github.com/lh3/seqtk, version 1.3; default parameter). 4) pWGS data mixed with a real background metagenomic dataset (WGSBG): Three different metagenomic datasets (see **Supplementary Table 4.** BG1: IBD, BG2: FMT, BG3: MI, as shown in **Figure 2**) were used to mix with the pWGS data of *E. coli* at different ratios (1, 5,10, 25 and 100-fold). Metagenomic samples were analyzed by Kraken2 (version 2.1.1; database: minikraken2_v2_8GB_201904) to ensure minimal abundance of *E. coli.* Strains of *E. coli* with pairwise genome wide ANI between 95%-99% were selected to represent different subspecies (**Supplementary Table 3**). In each synthetic data set of mixed strains, 20 combinations of strain composition were generated by Dirichlet distribution (**Supplementary Table 2**). All synthetic datasets were generated by the SimStr pipeline (https://github.com/xbiome/StrainPanDA/tree/main/SimStr). For each strain, its genome size was considered 1× sequencing depth and used to calculate the number of reads to generate. The minimum relative abundance (*i.e.,* frequency) of a strain was set as 5% and as one unit. For example, for E. coli synthetic data of 1× sequencing depth that we refer to in this study, the data size of the strain with 5% frequency was ~4.5 megabases (MB), while the total depth of each sample in this dataset was always 20x, and ~90MB in size (1x sequencing depth as a unit and 20 units in total). To evaluate the effect of sample size, 400 synthetic mixtures of 4 strains were generated by Dirichlet distribution. The synthetic data were separated into 10 runs (40 samples each) and further downsampled to 20, 15, 10 and 5 samples in each run.

#### Synthetic data of gut bacterial species

Synthetic data of 6 species, including *B. longum, C. difficile, E. coli, E. faecalis, F. prausnitzii* and *P. copri,* were generated separately (Sync6, pWGS at 5 × sequencing depth) (**Supplementary Table 2**). For each species, the relative abundances of 4 strains (5%, 10%, 25% and 60%) were permutated to generate 24 samples in total (**Supplementary Table 2**). All 6 species were in the pre-built databases of StrainPanDA and PStrain. Only *B. longum, E. coli* and *E. faecalis* were in the pre-built database of StrainEst, so the other species were excluded in the comparison to StrainEst (**Fig. 2C**).

##### Evaluation of predicted strain composition

StrainEst and PStrain were run with their pre-built database and default parameters (StrainEst: ftp://ftp.fmach.it/metagenomics/strainest/ref/; PStrain: https://github.com/wshuai294/PStrain). The strain compositional profiles predicted by different methods were evaluated and compared by SimStr. For the predicted strain composition shown in Figure 2A (stacked bar plots), strains with relative abundance below 0.01 were filtered and the remaining strains were sorted by their relative abundance (rescaled to 1) in decreasing order. After sorting, the predicted strains in the lower tail exceeding the number of simulated strains were grouped into “Extras”.

Two commonly used metrics were used to evaluate the performance of predicted strain composition of different methods:

1) Jensen-Shannon divergence^74^ (JSD): JSD between the predicted strain composition and actual strain composition is calculated by the distance function in phyloseq^75^ (R package) on the sorted relative abundance (in decreasing order). If the number of predicted strains is different from the actual number of strains, zeros were appended to the vector with lower dimension. The JSD is symmetric and is in the interval of [0,1]. It reflects the dissimilarity in compositional profiles of strains, *i.e.*, JSD = 0 represents an exact prediction.

2) Matthews correlation coefficient^76^ (MCC):

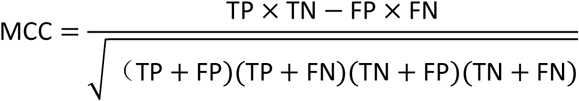

where TP is the number of true positives, TN is the number of true negatives, FP is the number of false positives, and FN is the number of false negatives. The MCC ranges from −1 to 1, where 1 represents an exact prediction, 0 represents a random prediction and −1 represents total disagreement.

Due to the lack of strain annotations from PStrain, we only computed MCC for strain composition predicted by StrainPanDA and StrainEst. For StrainEst, the predicted strains were directly annotated by reference genomes. For StrainPanDA, based on the predicted gene family profile, the Jaccard distance (JD(A, B) = *1* – |A ∩ B| ÷ |A ∪ B|) between the predicted strain and all reference genomes were calculated. The reference genome with the smallest Jaccard distance to the predicted strain was used for annotation. If a strain in the synthetic mixture is included in the pre-built database of reference genomes, ID of the annotated reference genome is directly compared to the actual strain to determine if the prediction strain is true positive. If a strain in the synthetic mixture is not included in the pre-built database of reference genomes, we used the phylogenetic tree to decide whether a predicted strain is true positive (**Supplementary Table 4**). Briefly, we generated a phylogenetic tree by parsnp^77^ (version 1.5.1, default parameter) including genomes of the strains used in synthetic mixtures and all the reference genomes. If the annotated reference genome of a predicted strain is within the cutoff of phylogenetic distance (cutoff = 0.05 for *E. coli,* corresponding to ANI ~99%) from an actual strain, it is considered as true positive.

##### Evaluation of predicted gene family profiles

For each microbial strain evaluated in benchmarking datasets, ErrFree reads at 5 × sequencing depth were generated by ART simulator^73^ (Sync-Single dataset) based on its reference genome downloaded from NCBI. The Sync-Single dataset had three replicates for each strain and was used to generate the actual gene family profile of each strain by PanPhlAn (default parameters). The gene families found in two or more replicates were considered as “present”. The actual gene family profile (Ground Truth) of each strain was compared to the predicted gene family profile (**Fig. 2D**). The Jaccard distances between the predicted gene family profiles of microbial strains and their ground truth profiles (or the gene family profile of a randomly sampled reference genome) were computed (**Supplementary Fig. 5**). The Precision-Recall curve of gene family profiles for each strain was generated by R package PRROC^78^ using the confidence scores to rank the gene families predicted (**Supplementary Fig. 6**). For random guesses, 1000 random gene family profiles were generated by sampling *N* gene families from the pangenome as “present”, where *N* is the average number of gene families present in reference genomes. To demonstrate the ability of StrainPanDA to identify strain-specific genes, the pathogenic *E. coli* strain *O*104 (GCF_002983645) was introduced in a synthetic dataset of 4 strains (Sync *O*104 dataset, pWGS, 5× sequencing depth). The outbreak related genes curated from Segata *et al.* ^35^ were used to evaluate StrainPanDA’s gene content prediction (**Supplementary Table 5**). We also compared the predicted gene family profiles from StrainPanDA to the prediction from PanPhlAn (**Supplementary Fig. 8)**. To simplify the output, only samples with the highest abundance of one strain were used for PanPhlAn prediction.

### Applications of StrainPanDA in metagenomic data

#### Case study: mother-infant gut metagenomes

All available samples of ERP005989 were downloaded from EBI (8 samples failed, **Supplementary Table 6)**. Based on diet history, infants without diet history were filtered. The rest of 84 infants were split into 3 different groups (B_F_F: discontinued breastfeeding at 4-month; B_B_F: discontinued breastfeeding at 12-month; B_B_B: continued breastfeeding; the F_B_M sample was excluded) (**Supplementary Table 6**). Samples without enough coverage on *B. longum* gene families were filtered by StrainPanDA at the preprocessing step and excluded from the downstream analysis. In the “continued breastfeeding” group, infants that kept breastfeeding between successive time points (*i.e.*, between newborn and 4-month, or between 4-month and 12-month) were included. In the “discontinued breastfeeding” group, infants which stopped breastfeeding by 4-month or 12-month were included. For functional interpretation of the subspecies, we grouped the gene families annotated to the same KEGG ortholog or CAZy family. To further analyze the key functions related to breastfeeding, we curated a set of KEGG orthologs from related references^51,79^. All the KEGG orthologs were further grouped into subclasses and classes based on the literature^51,79^ (**Supplementary Table 7**). The gene family coverage was calculated as the fraction of detected genes belonging to the subclass (**Fig. 3C**).

#### Case study: FMT donor-recipient metagenomes

Raw sequencing reads were downloaded from ENA (Accession: PRJNA625520 for the study on Crohn’s disease^37^, PRJEB23524 for the study on *C. difficile* infection study^29^). For the Crohn’s disease dataset, species relative abundances were estimated using Kraken2^18^ (version 2.0.8-beta) with the miniKraken database (v2_8GB_201904_UPDATE). To identify species associated with remission or relapse, Wilcoxon rank sum test was conducted to select differentially abundant species using the mean relative abundances across different timepoints for each subject (samples collected before FMT or after relapse were discarded). The relative abundances of subspecies were predicted by StrainPanDA and normalized by the centered-log-ratio transformation for calculating Spearman correlation with the species abundances. For functional annotation of the subspecies, virulence factors and CAZy annotations were taken from the species-specific databases constructed as described above.

##### Functional annotation of gene families

To annotate the gene families by KEGG, gene family representative sequences were mapped against KEGG orthologs (release 2020-07-20) using usearch^69^ (v11.0.667; ‘-ublast’). Alignments with identity > 50% and query coverage > 50% are kept. To annotate the gene families by CAZy, seqkit^80^ (0.15.0) was used to translate the gene family centroids into six open reading frames. The translated amino acid was used as the input of run_dbcan^81^ (2.0.11), which used DIAMOND^82^ (2.0.8), HMMer^83^ (3.3.2) and Hotpep^84^ (2.0.8) with default parameters to predict the CAZy annotation. The CAZy annotations was selected only if it was predicted by at least two programs. If a gene family was assigned by more than one CAZy annotations, only the first annotation was used. To annotate virulence factors, gene family centroids were mapped against the Virulence Factor Database (VFDB, Apr 9 2021) by DIAMOND^82^ (2.0.8) (blastp, query coverage >50% and identity > 50%).

## Declarations

## Acknowledgements

We would like to thank Haokui Zhou, Ramnik Xavier and members of Lei Dai lab for constructive comments on the manuscript. We would like to thank Jinhui Tang and Yong Liang for their assistance in bioinformatics analysis, Xi Wang and Dongdong Xu for their support in maintaining the computing environment, Reese Hitchings for proofreading the manuscript.

## Software availability

The source code of StrainPanDA is available on Github (https://github.com/xbiome/StrainPanDA).

## Authors’ contributions

H.H., Y.T. and L.D. conceived and supervised the study. H.H., Y-X.T. and C.L. developed the algorithm and performed the analysis on simulated and real metagenomic samples. L.D., H.H., Y-X.T. and C.L. wrote the manuscript with inputs from all co-authors.

## Ethics approval and consent to participate

Not applicable

## Consent for publication

Not applicable

