## Supplementary Figures for "StrainPanDA: linked reconstruction of strain composition and gene content profiles via pangenome-based decomposition of metagenomic data"

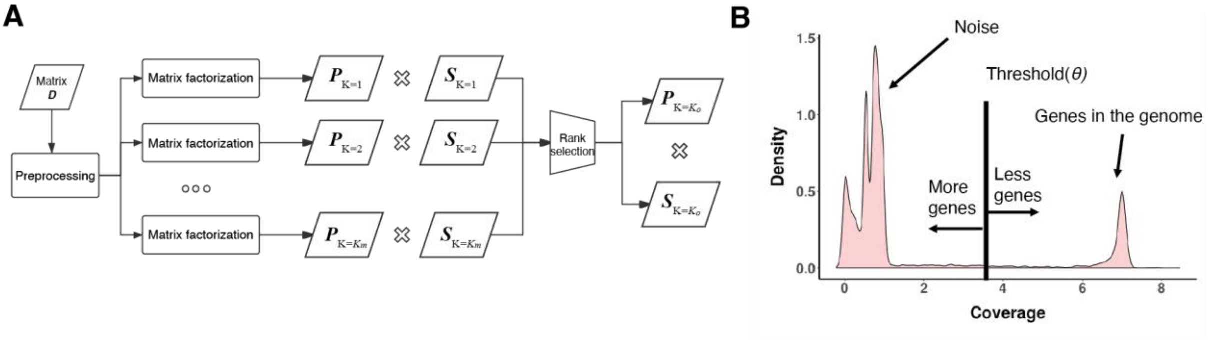


**Supplementary Figure 1. Details of StrainPanDA algorithm. (A)** Illustration of the determination of strain number (*i.e.*, factorization rank) in non-negative matrix factorization. The gene family abundance matrix ***D*** was decomposed into the product of a gene content profile matrix ***P*** and a strain composition matrix ***S*** at different number of strains (K=1 to $K_m$, $K_m$=15 by default). The rank selection step chooses a proper K (see Methods for details). **(B)** Illustration of the determination of gene family presence (***P*** matrix). A threshold (vertical line) is used to determine the presence/absence of the gene families and confidence scores are calculated as described in Methods.

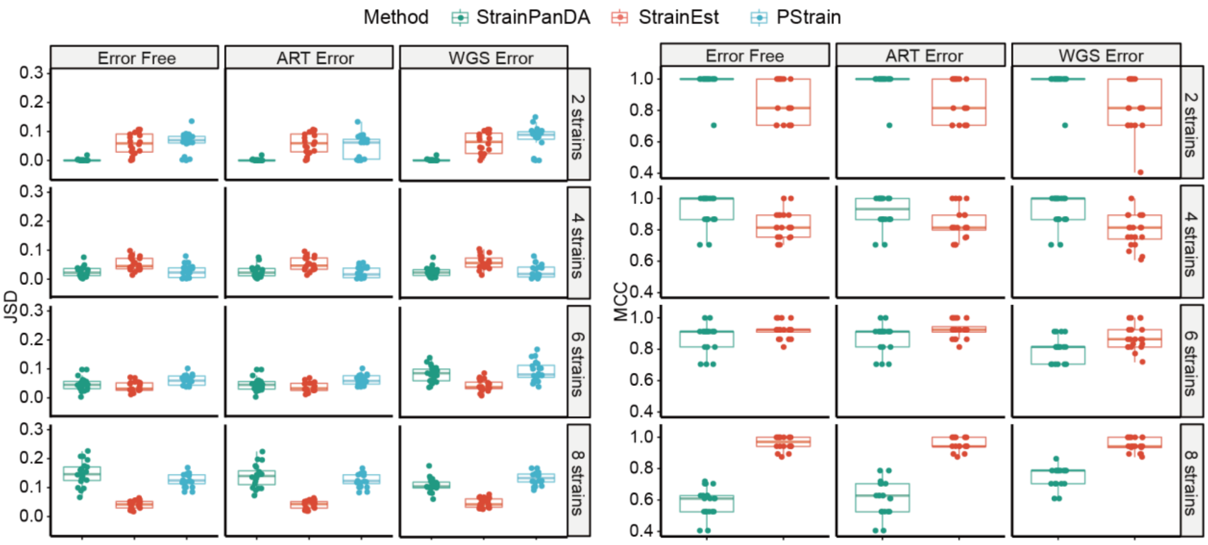


**Supplementary Figure 2. Comparison of JSD and MCC of predicted *E. coli* strain compositions among StrainPanDA, StrainEst, and PStrain using error free data (Error Free), data with sequencing error simulated using ART software (ART Error), and real sequencing data for the corresponding ground truth isolate (WGS Error, or "pWGS") at 1× sequencing depth.**

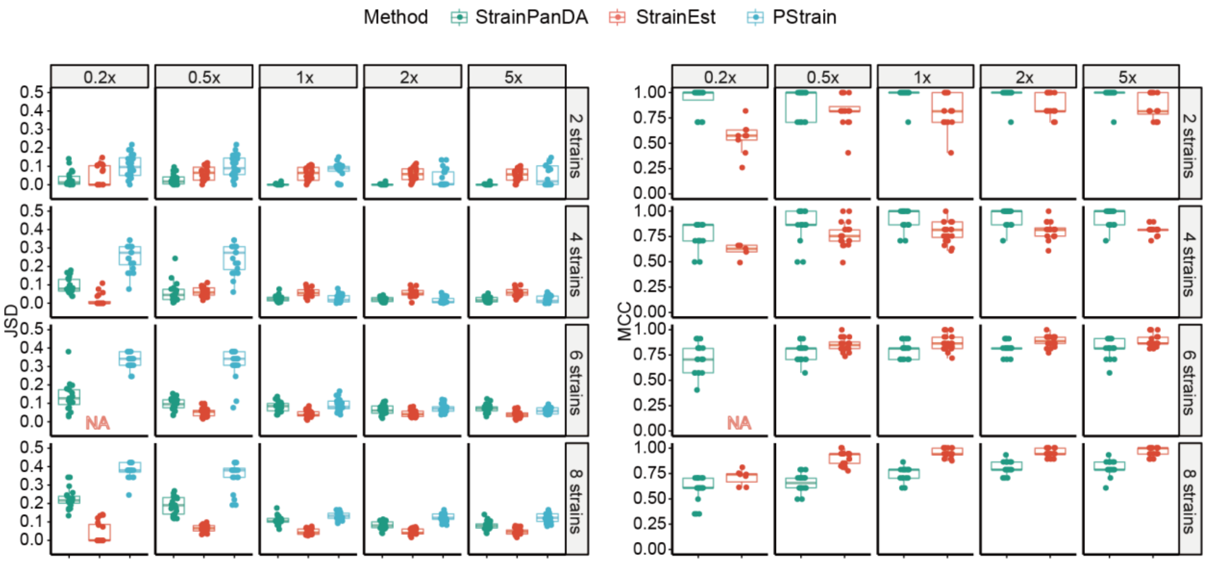


**Supplementary Figure 3 Comparison of JSD and MCC of predicted *E. coli* strain compositions among StrainPanDA, StrainEst, and PStrain using pWGS datasets of 0.2×, 0.5×, 1×, 2× and 5× sequencing depth.** Output not available is marked as "NA". MCC was not available for PStrain due to its lack of strain annotation.

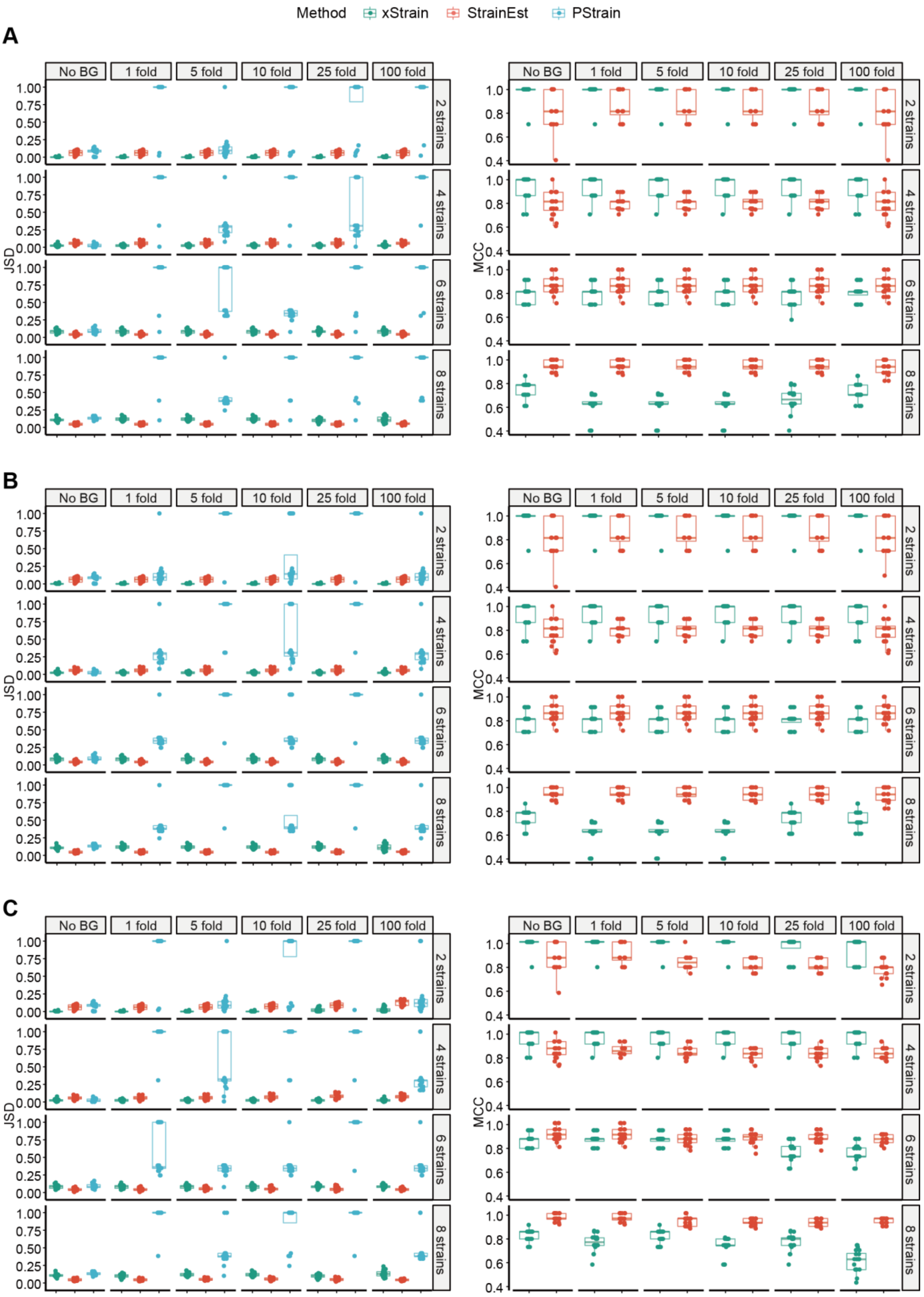


**Supplementary Figure 4. Comparison of JSD and MCC of predicted *E. coli* strain compositions among StrainPanDA, StrainEst, and PStrain on datasets of 1× sequencing depth and spike-in *E.***

***coli* WGS reads with 1-fold, 5-fold, 10-fold, 25-fold and 100-fold background reads from the IBD (panel A) (https://ibdmdb.org/), FMT (panel B) (Smillie *et al.* 2018) and MI (panel C) (Bäckhed *et al.* 2015) datasets.** MCC was not available for PStrain due to its lack of strain annotations.

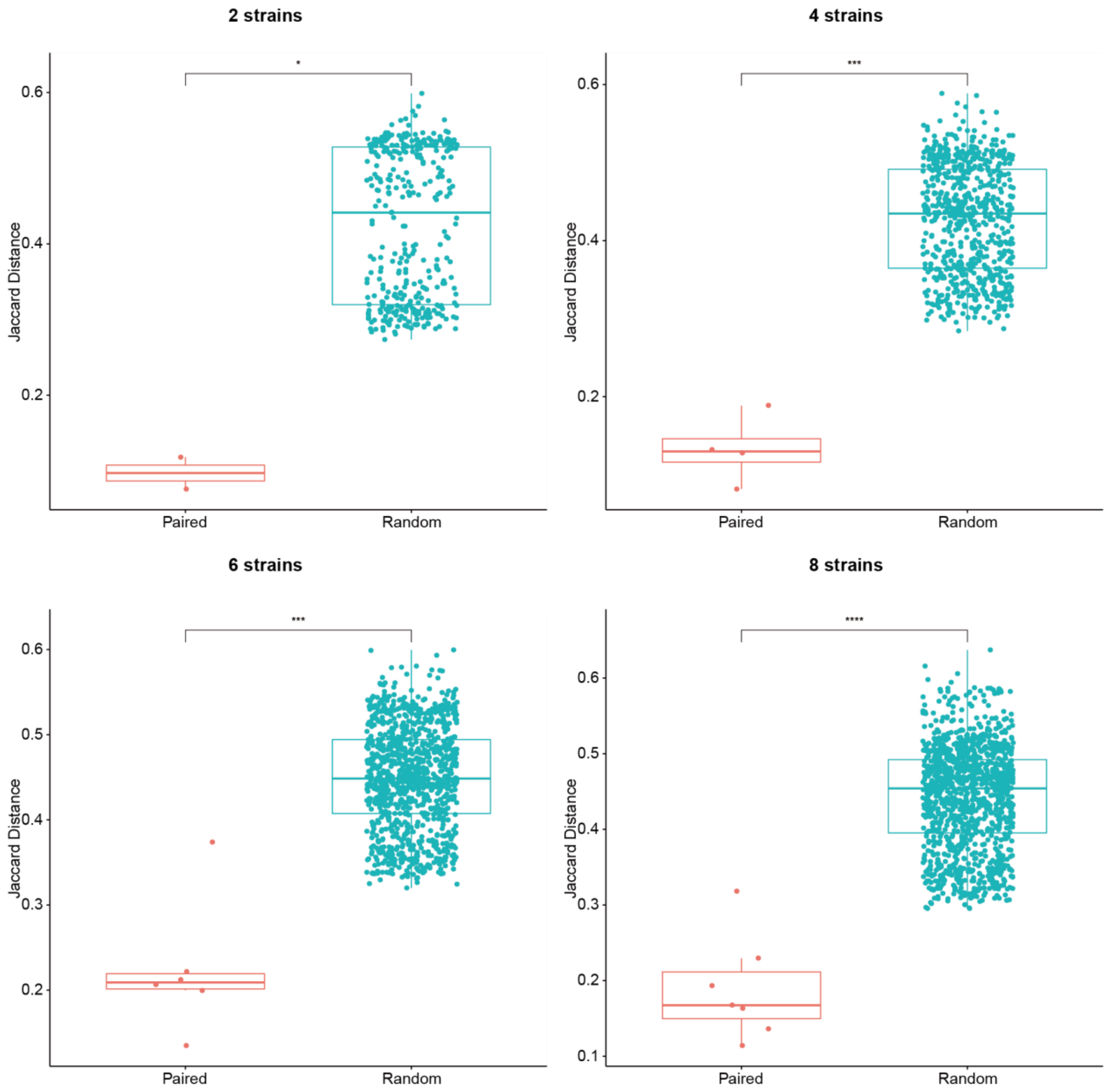


**Supplementary Figure 5**. **Jaccard distance between the predicted gene family profile of *E. coli* strains and the ground truth (Paired) is much smaller than the distance to randomly sampled reference genomes (Random)**. pWGS datasets at 1× sequencing depth with different strain numbers (2, 4, 6 and 8 strains). P values from t-test; *P < 0.05, ***P < 0.001, ****P < 0.0001.

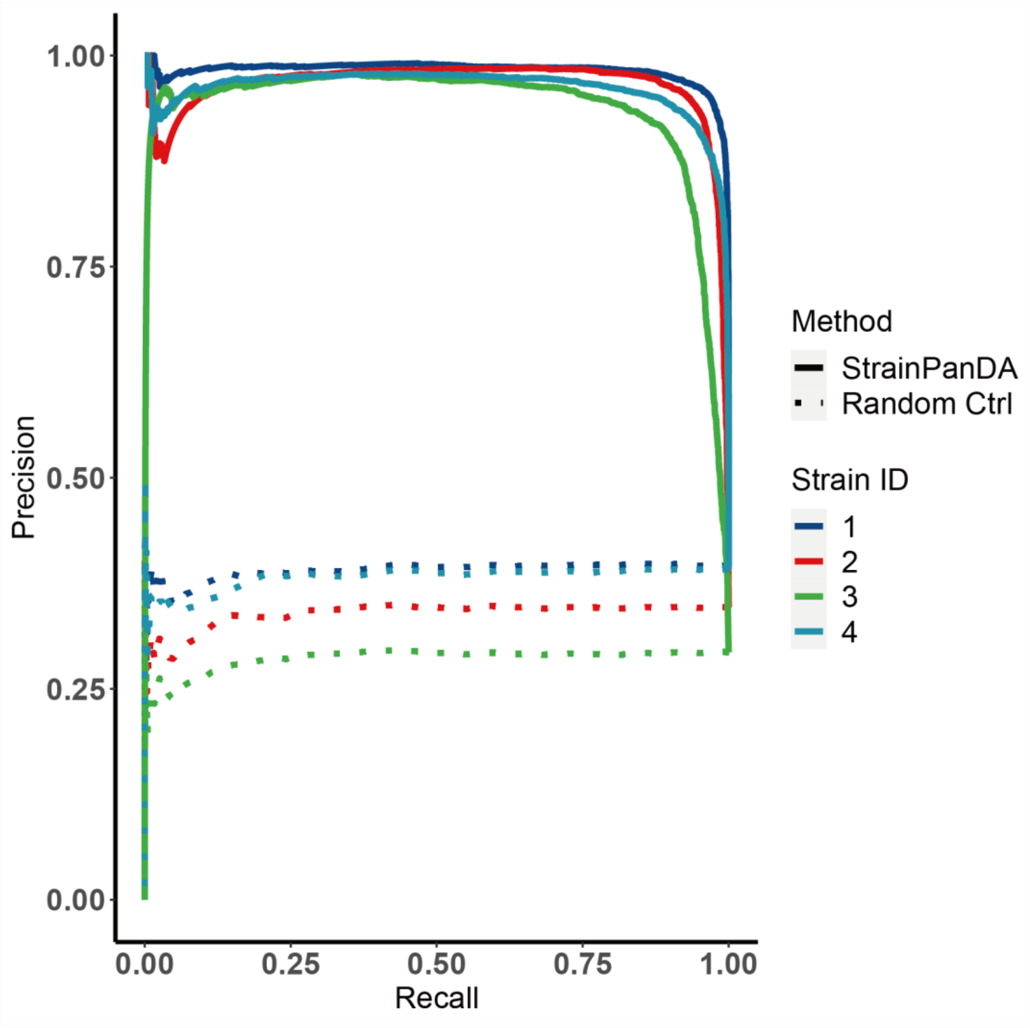


**Supplementary Figure 6. Precision-Recall curve of predicted gene family profiles of *E. coli* strains.** Solid line: prediction by StrainPanDA; dotted line: a randomly generated gene family profile as the control. Synthetic mixtures of *E. coli* strains: pWGS dataset, 1×sequencing depth.

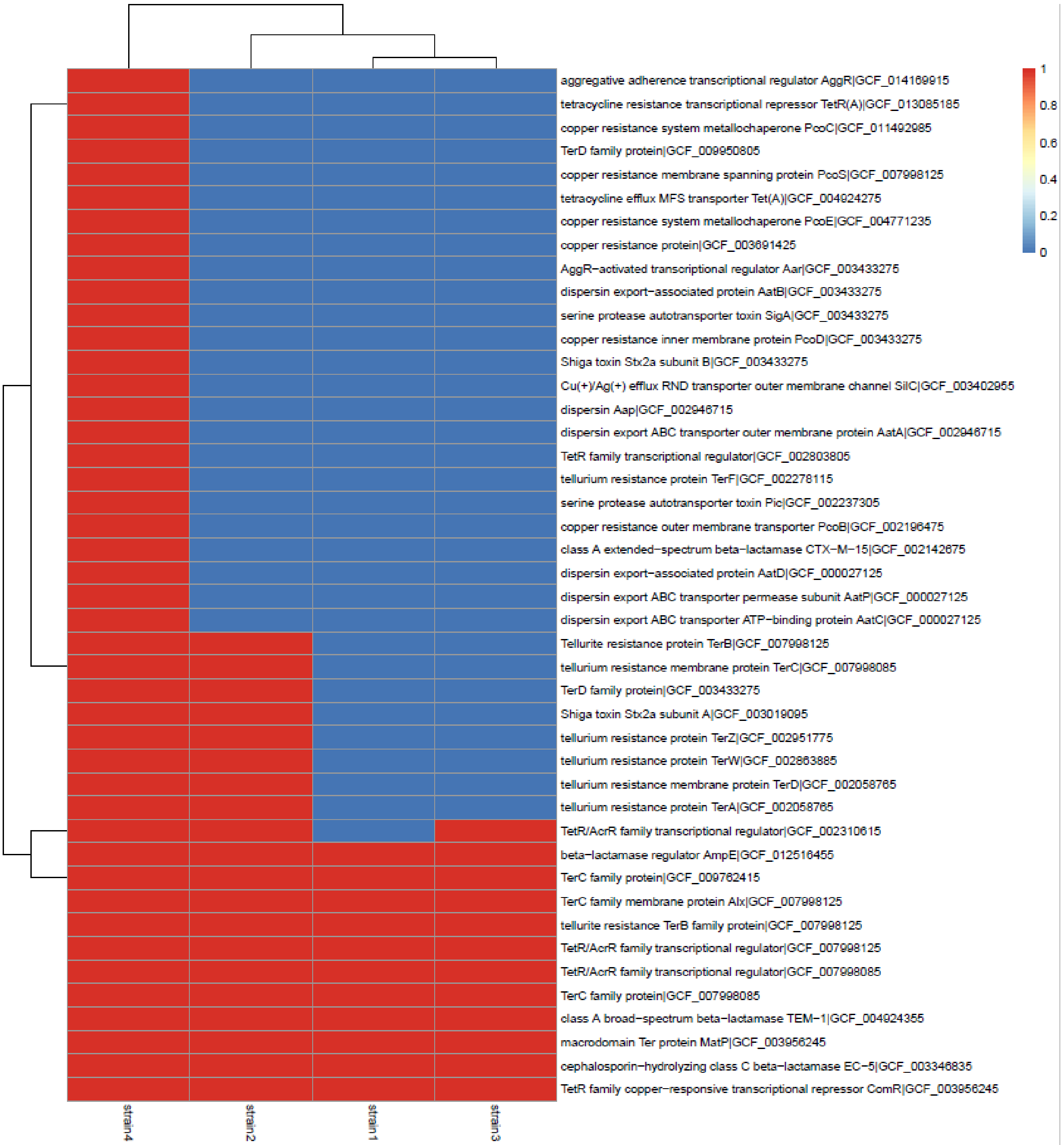


**Supplementary Figure 7. Identification of *E. coli* strain-specific outbreak-related gene families by StrainPanDA.** Red/blue: presence/absence of a gene family. Strain4 is the *E. coli* O104 strain in synthetic mixture (Sync-O104 dataset).

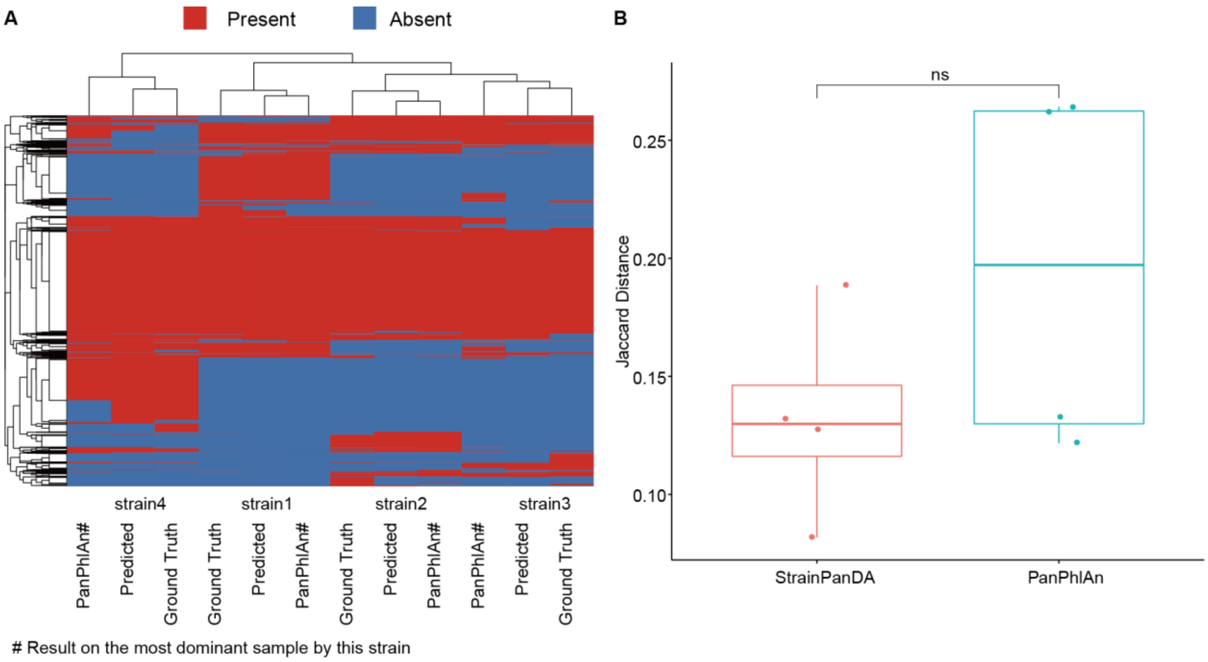


**Supplementary Figure 8. Predicted gene content profile of the dominant strain in benchmarking samples by StrainPanDA and PanPhlAn. (A)** The ground truth and the predicted gene family profiles by StrainPanDA and PanPhlAn of *E. coli* strains (pWGS dataset, 1× sequencing depth). Each row is one gene family, and each column is one strain. Hierarchical clustering on rows and columns are performed based on Euclidean distance. **(B)** Jaccard distance between the predicted gene content profiles and the ground truth (n = 4 strains). ns: not significant, paired t-test.

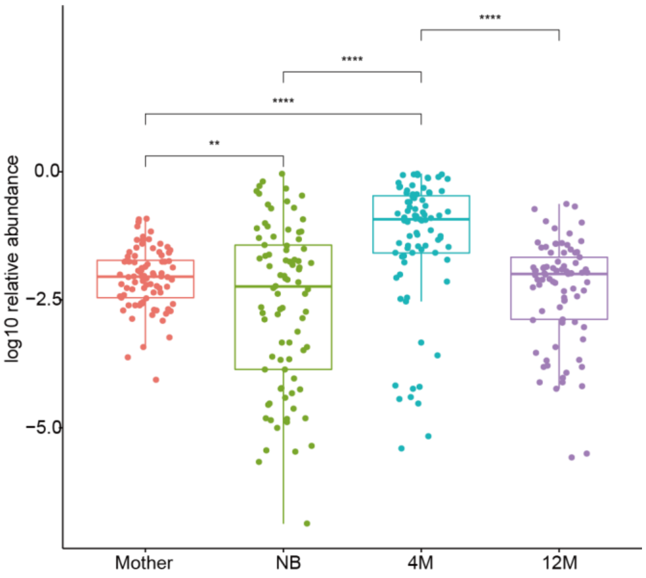


**Supplementary Figure 9. Relative abundance of *B. longum* in the gut microbiome of mothers and infants (3 different time points).** NB: newborn, 4M: 4-month, 12M: 12-month. P values from t-test: **P < 0.01, ****P < 0.0001.

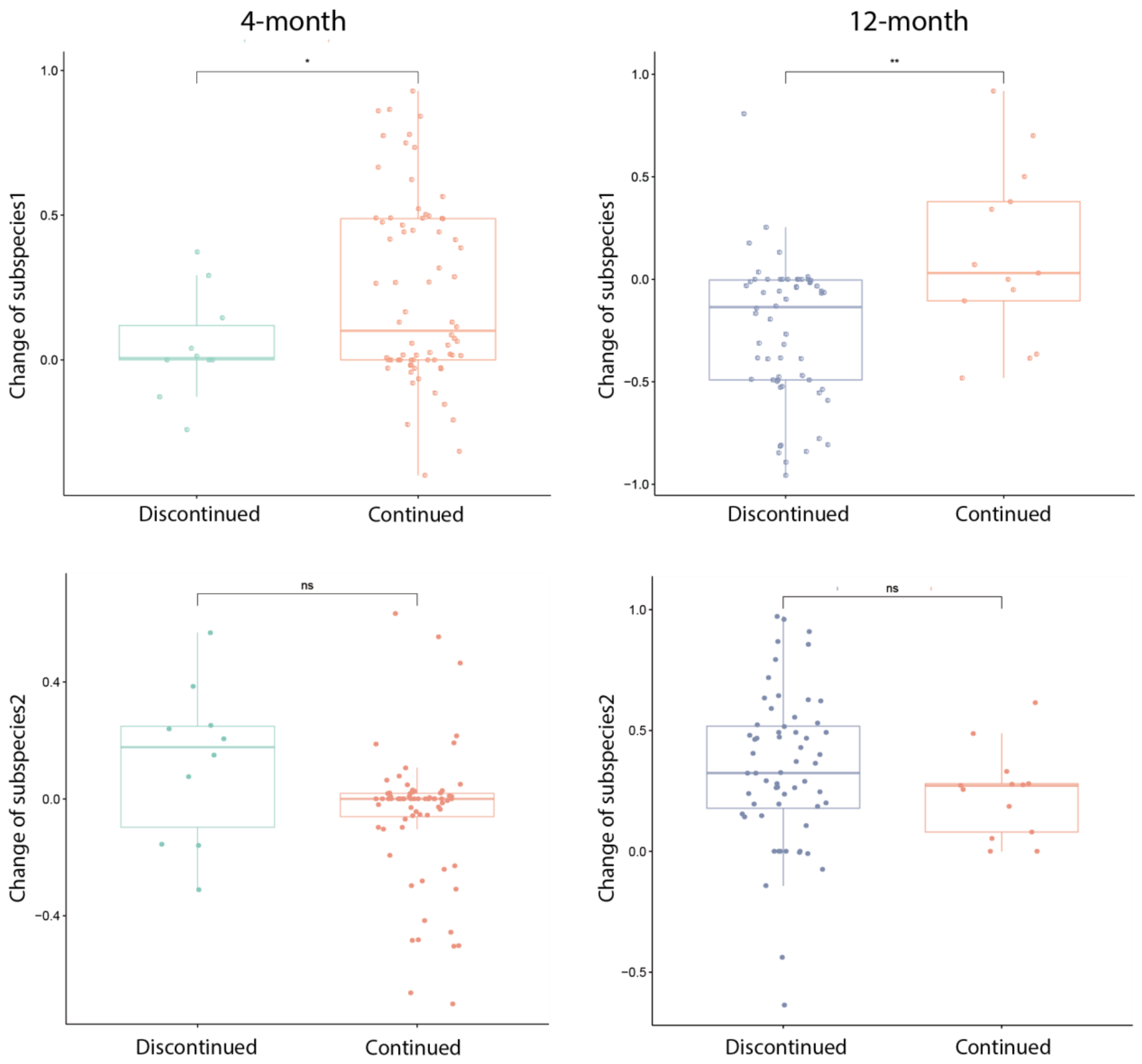


**Supplementary Figure 10. Difference in the relative abundance of *B. longum* subspecies 1 and subspecies 2 of infant gut microbiome between successive time points.** Red: continued breastfeeding; green: discontinued at 4 months; purple: discontinued at 12 months. P values from t-test: *P<0.05, **P < 0.01, ns: not significant.

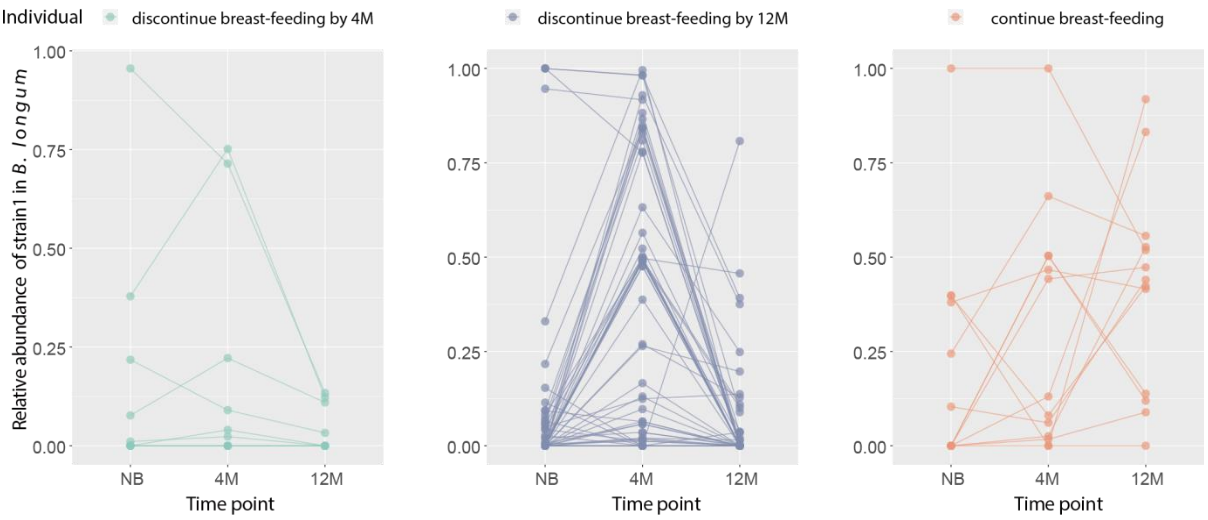


**Supplementary Figure 11. The dynamics of *B. longum* subspecies 1 in infant gut microbiome is affected by breastfeeding.** NB: newborn, 4M: 4-month, 12M: 12-month.

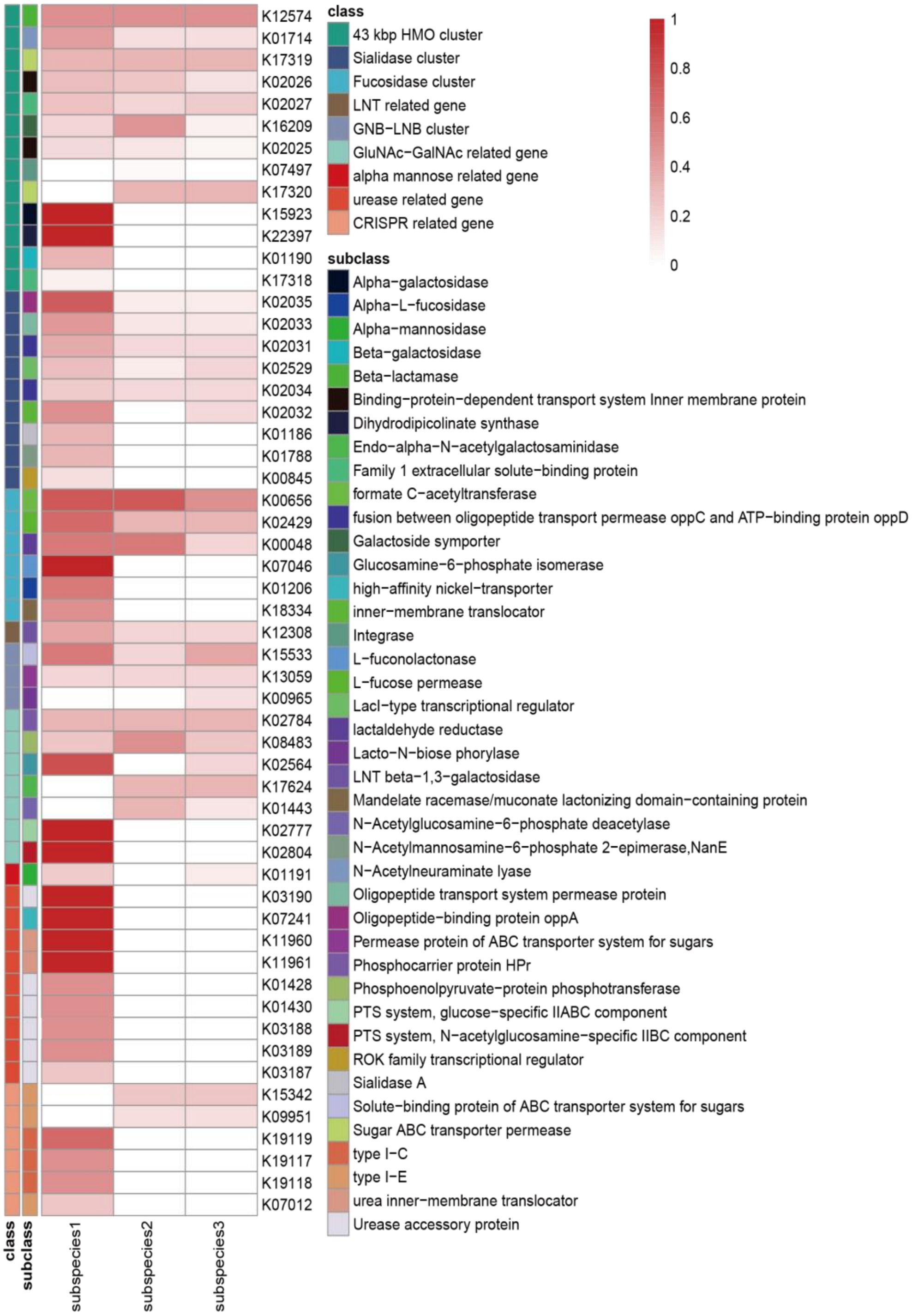


**Supplementary Figure 12. Functional variations of predicted *B. longum* subspecies.** Gene families related to the metabolism of host glycans, urease and CRISPR are annotated by with the KEGG database. Columns are subspecies and rows are KO ID of genes. The color scale in heatmap indicates the normalized coverage of gene families of the specific KO ID.

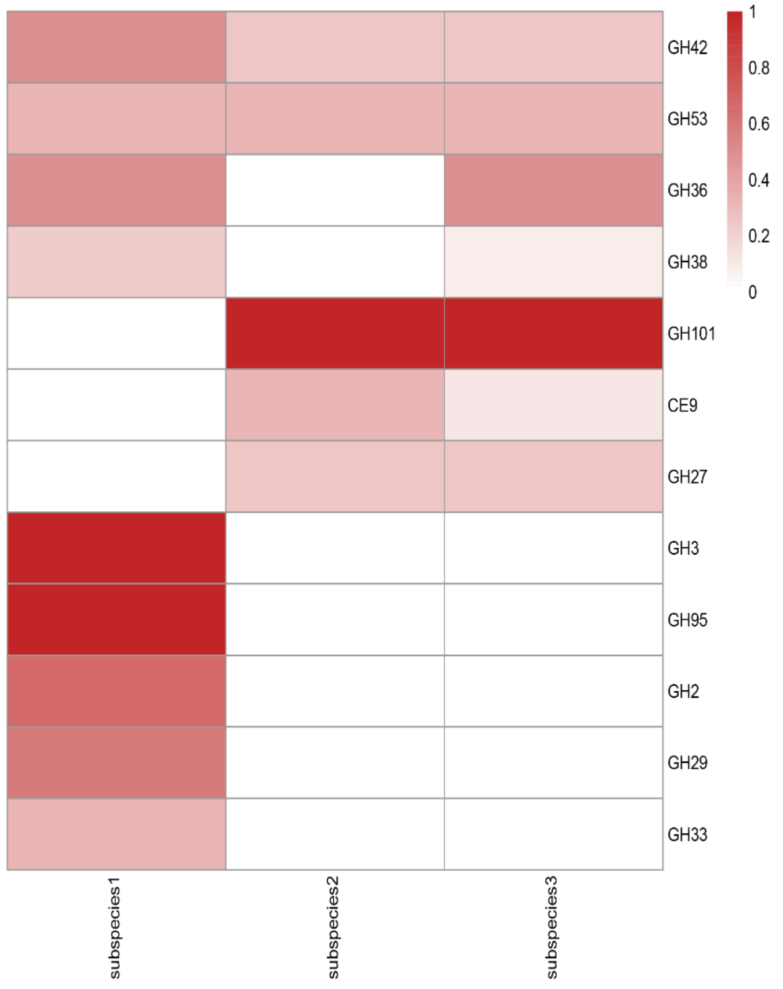


**Supplementary Figure 13. Heatmap of functional variations of predicted *B. longum* subspecies on genes related to the metabolism of host glycans by CAZy annotations.** Columns are subspecies and rows are CAZy families. The color scale in heatmap indicates the normalized gene coverage in the specific CAZy family.

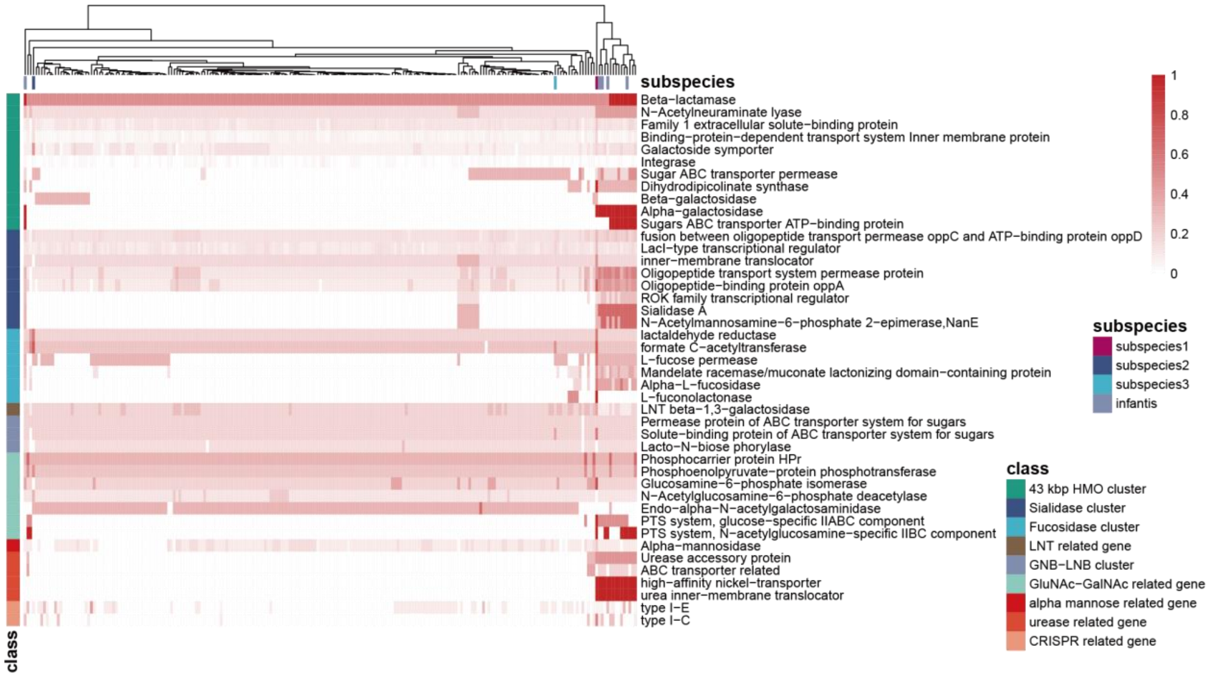


**Supplementary Figure 14. Clustering of predicted subspecies and reference genomes of *B. longum*.** Each column was a strain/subspecies, and each row was a subclass of function. The color scale in heatmap indicates the normalized gene coverage in the specific subclass (i.e. the fraction of detected genes belonging to the subclass). Strains previously annotated as *B. longum* subspecies *infantis* were highlighted.

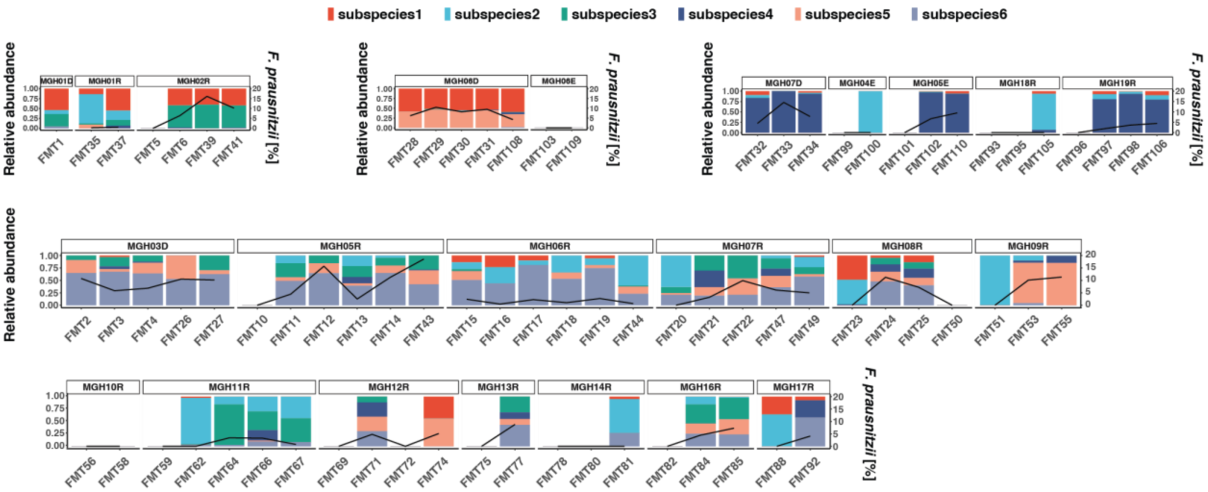


**Supplementary Figure 15. StrainPanDA analysis of *F. prausnitzii* in a metagenomic dataset of *Clostridium difficile* infected patients treated by fecal microbiota transplantation (FMT).**

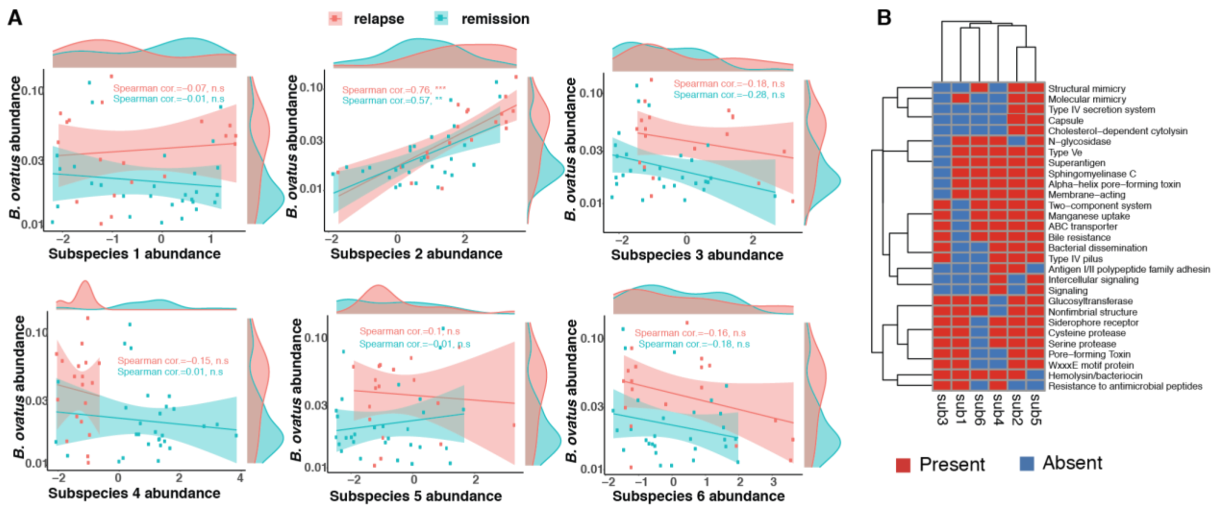


**Supplementary Figure 16. StrainPanDA analysis of *B. ovatus* in a metagenomic dataset of Crohn's disease patients treated by fecal microbiota transplantation (FMT). (A**) Scatter plots with fitted linear regression lines (with 95% confidence interval represented by the shaded areas) showing the relationship between the abundances of *B. ovatus* and its subspecies (normalized by centered log-ratio transformation). The density plot on the side of each panel shows the marginal distribution of the corresponding variable. **(B)** Virulence factor profile of the *B. ovatus* subspecies (virulence factors shared by all the subspecies were not shown).

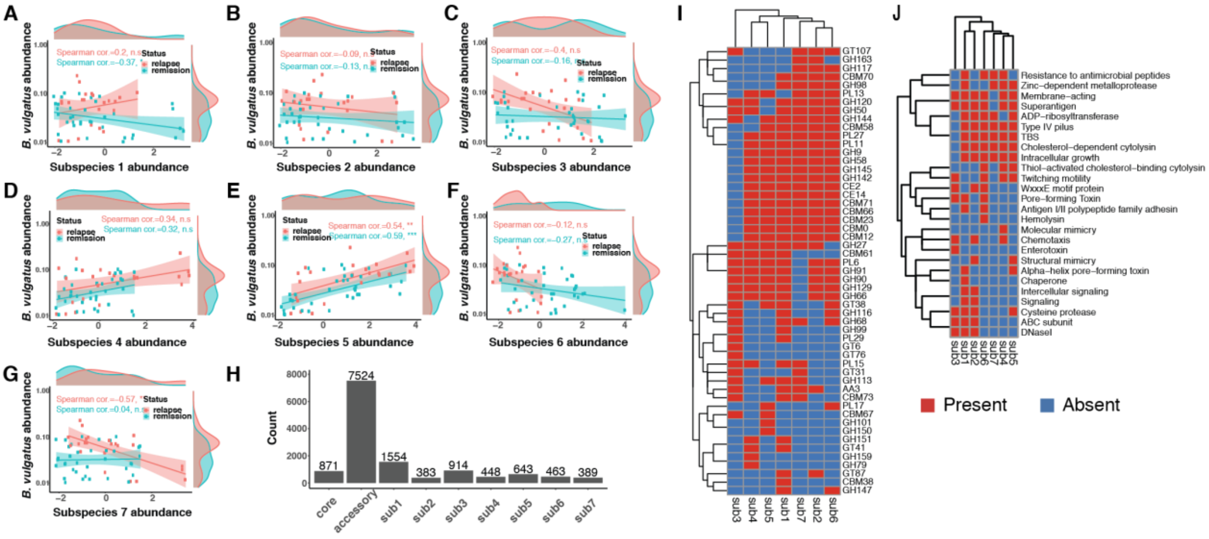

**Supplementary Figure 17. StrainPanDA analysis of *B. vulgatus* in a metagenomic dataset of Crohn's disease patients treated by fecal microbiota transplantation (FMT). (A-G)** Scatter plots with fitted linear regression lines (with 95% confidence interval represented by the shaded areas) showing the relationship between the abundances of *B. vulgatus* and its subspecies (normalized by centered log-ratio transformation). The density plot on the side of each panel shows the marginal distribution of the corresponding variable. **(H)** Barplot showing the summarized pangenome information of *B. vulgatus* present in the dataset. **(I)** Carbohydrate-Active enZYmes (CAZy) profile of the *B. vulgatus* subspecies (CAZy families shared by all the strains were omitted). **(J)** Virulence factor profile of the *B. vulgatus* subspecies (virulence factors shared by all the subspecies were not shown).

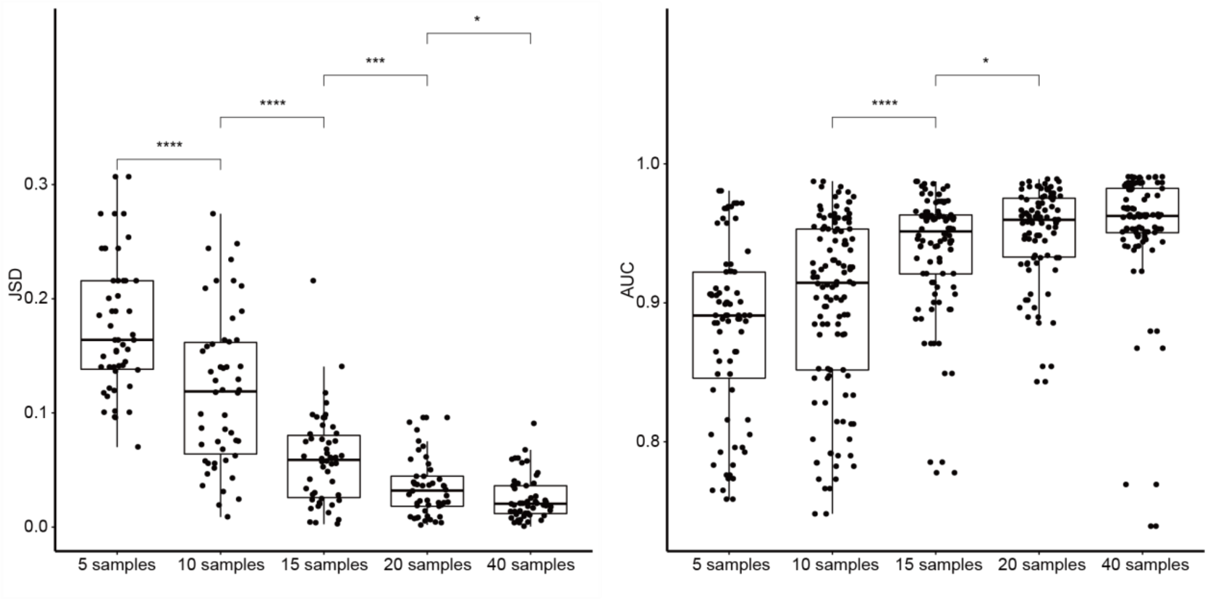


**Supplementary Figure 18. Evaluation of StrainPanDA using synthetic data with varying sample size.** JSD quantifies the accuracy of predicted strain composition, while AUPRC quantifies the accuracy of predicted gene content profile of strains. P values from t-test: *P < 0.05, ***P < 0.001, ****P < 0.0001.
